# Quantitative genetics of natural *S. cerevisiae* strains upon sexual mating reveals heritable determinants of cellular fitness

**DOI:** 10.1101/2024.02.12.579867

**Authors:** Sivan Kaminski Strauss, Ruthie Golomb, Dayag Sheykhkarimli, Gianni Liti, Orna Dahan, Yitzhak Pilpel

**Affiliations:** Department of Genetics, Weizmann Institute of Science. Rehovot, 76100, Israel; CNRS, INSERM, IRCAN, Côte d’Azur University, Nice, France; Donnelly Centre and Departments of Molecular Genetics and Computer Science, University of Toronto, Toronto, ON, Canada; Lunenfeld-Tanenbaum Research Institute, Sinai Health System, Toronto, ON, Canada

## Abstract

Quantitative genetics requires large datasets of diverse phenotyped-genotyped strains from the same species. A special need is for such archived biological material and computerized data in sexually reproducing individuals from a species. Here we leverage sexual mating among close to 100 diverse natural isolates of the yeast *S. cerevisiae* that form about 4,000 hybrids combinations in several ecologically relevant growth conditions. In a first genetic study of this new resource we focus on fitness measurements and its modes of inheritance as a quantitative trait from parents to offspring hybrids. We employ genomic barcoding of all strains and a barcode recombination technique to follow hybrids of each successful mate combination. For all parents, and separately for all offspring hybrids we measure fitness under each condition. We focus on the inheritance of fitness, the ultimate evolutionary trait, and its inheritance as a quantitative trait upon sexual mating. Predicting hybrid fitness given parental parameters is a major challenge as it is likely multi-factorial. We find that hybrids fitness in fermentable carbon source correlates positively, yet modestly, with parental fitness, while on non-fermentable carbon, hybrid fitness shows no detectable correlation with parental fitness. Instead, the non-fermentable condition, hybrid fitness increases sharply with genetic distance between their parents, suggesting that outbreeding maximizes fitness irrespective of parental fitness at that condition. The number of minor alleles in the genome of each hybrid, analogous to polygenic risk score in classical genetics, negatively correlates with fitness in both conditions. Fitness inheritance can be explained by either a dominance or a co-dominance modes of inheritance, in the non-fermentable and fermentable conditions respectively. Our newly suggested biological resource and data provide new foundations for a quantitative research in genetics and evolution upon sexual mating. Furthermore, our barcoded strains and mating tracking method provide an important research resource for the yeast community.

## Introduction

A strong quantitative basis for genetics requires extensive dataset of genotyped and phenotyped individuals from a species. A significant distinction is between genetics of asexual-vegetative and sexual life modes. Sexual reproduction introduces considerable complexity in the research, especially with regards to the inheritance patterns of quantitative traits. Quantifying the inheritance of such traits upon sexual mating involves elucidating and predicting trait values among offspring based on various parental parameters. A pivotal quantitative trait in evolutionary dynamics is an organism’s fitness, encompassing survival and reproductive rates in diverse environments [1]. Fitness is determined by an intricate interplay of various cellular and molecular traits and parameters. Gene expression is one such key trait that significantly impacts fitness [2]. Unraveling the inheritance of fitness and of gene expression following sexual mating presents a significant challenge for modern genetics. Deciphering the modes of inheritance of these traits upon sexual mating can also provide an important conceptual means to understand the impacts of sex on evolution.

### Sexual mating in yeast

The budding yeast, *Saccharomyces cerevisiae*, is widely used as a model system for cell biology, genomics and genetics research [3]–[6]. In recent years, the number of sequenced genomes of strains from this and related species and other genetic tools in yeast are increasing [7]–[17]. Yeast is also used for studying the interaction between genotype and phenotype (e.g., genome wide association studies (GWAS) [9], [15]–[17] and quantitative trait loci (QTL) studies [13], [14], [18]). The 1011 natural yeast isolates study that was published by Peter *et al* added many different genomes, as well as their genomic and phenotypic characteristic as a tool for studying population genomics and yeast phylogenetic [9]. The budding yeast *S. cerevisiae*, can grow either vegetatively as haploid or as diploid, and can also mate sexually. Following pheromone sensing, haploid cells (i.e., Mat**a** or Matα) mate to generate a diploid cell that can then continue growing vegetatively (refer to here as “hybrid”). Upon nitrogen and carbon starvation, diploid cells undergo sporulation and meiosis to form haploid spores. At the end of the process 4 spores are produced and are stored within the intact cytoplasm of the mother cells, the ascus [19].

Most traits show parent-offspring correlation, in which offspring take after their parents [17]. To study trait inheritance, a large sample of parents and offspring should be used and analyzed [17], [18] which might be difficult to obtain in nature, or in many model organisms in the lab. The budding yeast is ideal for overcoming these challenges; first, haploid yeast cells can be easily crossed to achieve many diploid hybrids. Second, as yeast are small and easy to grow in the lab, one can produce and study as many as 10E8 cells (which in a single cell organism such as yeast translate to 10E8 organisms) in a single test tube in a single day. Third, yeast are easily manipulated genetically, and indeed many studies have used it to produce different libraries [20]–[22]. Thus, yeast have the potential to be used as a tool for the study of trait inheritance in a high-throughput manner.

### Fitness inheritance

Fitness is multi-genic, and involve many loci in the genome. As other quantitative traits, fitness of offspring is likely to be dependent upon parental fitness. In addition, fitness is known to be affected by the genetic distance between parents, which in extreme cases is responsible for two phenomena, heterosis, on the one hand, and genetic lethality on the other [23]–[31].

<u>Heterosis,</u> or hybrid vigor, is defined as the increased function of hybrid fitness compared to its parents. A hybrid is heterotic if its traits are enhanced as a result of mixing the genetic contributions of its parents. The heterotic offspring often has traits that are more than the simple addition of the parents’ traits, and can be explained by Mendelian or non-Mendelian inheritance. Typical heterotic/hybrid traits of interest in agriculture are higher yield, quicker maturity, stability, drought tolerance etc. [32].

In this work, we have leveraged on Peter *et al*. natural isolates collection [9] to create a platform for the study of different trait inheritance. We have a created a library with ∼100 parents haploid strain, each contains a combination of two barcodes that can be fused in the hybrid [33] (and similarly to [14]). Antibiotic resistance markers as well as fluorescence markers were also added to parental strains. We present the use of this platform to study fitness inheritance in different environments. We measure the fitness of ∼100 parents haploid, as well as their 4000 hybrids, and characterize their parent-hybrid fitness correlation as well as the effect of parental genetic distance on hybrid fitness. We find limited yet statistically significant inheritance of fitness between pairs of parental strains and their hybrid. Yet, this dependency of hybrid fitness on their parents is very conditions dependent. Further, hybrid fitness was found to be a function of the genetic distance between parents, especially in non-fermentative conditions where it is maximized upon outbreeding of relatively genetically distant strains.

## Results

### A massively parallel barcode recombination system for tracking yeast hybrids and their fitness

To study fitness inheritance of quantitative traits upon sexual mating in yeast we have generated a collection of genomically barcoded and labeled natural strains based on the *S. cerevisiae* natural strains [9]. From the original collection, ∼100 strains were chosen that vary and span a broad range of growth fitness and pairwise genetic distance (GD) (**Figure S1** and see **supplementary note 1**). To allow high-throughput experiments, these natural strains were genetically engineered to be used as haploid parental strains and as such, to allow detection, rate of formation, and relative fitness of all their hybrids combinations. In short, the HO locus [34], [35] was knocked out by a construct containing an antibiotic resistance cassettes, a fluorescence marker, and a barcode recombination element, based on the Cre-lox recombinase (barcode fusion genetics, BFG) [33] (and similarly to [36]) (**Figure S2A, supplementary note 1**). This cassette allows recombination of haploid parental unique barcodes (from the two mating partners, Mat**a** and Matα) into a fused barcode on the same DNA segment that can be sequenced (**Figure 1A, Figure S2, supplementary note 1**). To perform high-throughput experiments, all parental haploid strains from the two mating types were mixed and allowed to mate *en masse* (**Figure 1B**). After mating, diploid cells were retained and haploids that did not mate were eliminated (see Materials & Methods). Hybrids diploids (as well as Mat**a** and Matα parental haploids) were subjected to a competition assay to measure their relative fitness [37], [38] (**Figure 1C**).

**Figure 1.**
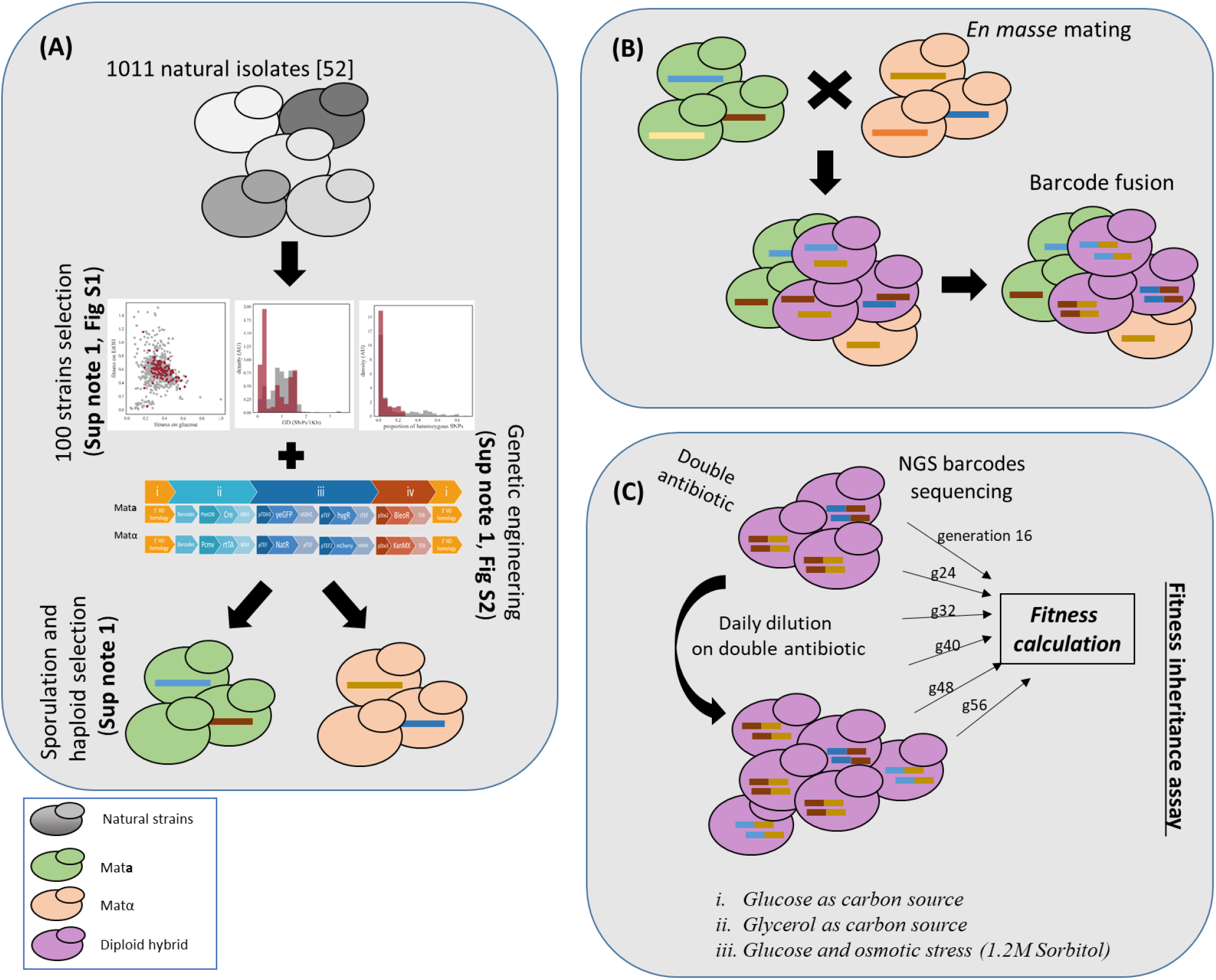
The experimental scheme to trace all mate pairs and fitness inheritance. (A) ∼100 strains from Peter *et al*. [9] 1011 strains were chosen according to three parameters; their fitness under respiration and fermentation conditions, the entire set GD, and heterozygosity levels (see **supplementary note 1** and **Figure S1**). The 100 strains were genetically manipulated to contain fluorescence and antibiotic markers and a barcode fusion genetic system (see **supplementary note 1** and **Figure S2**). Mat**a** and Matα final strains were achieved by sporulation and haploid selection (see Materials & Methods, and **supplementary note 1)** (B)All strains from both mating types (Mat**a** and Matα) were mixed together and allowed to mate with one another, *en masse*. BFG was enabled during mating and throughout the first 8 generations of vegetative growth (in the competition experiments Figure 1C). (C)Fitness inheritances assay: hybrids diploids were selected using two antibiotics to allow hybrids growth only. Cultures were grown for ∼60 generations in a daily dilution, and were sequenced in 6 time points, from generation 16 and then every 8 generations (see Materials & Methods). Fitness was calculated using a Maximum-Likelihood (ML) algorithm [37], [38].

### General characteristics of parental and hybrids strains

The strain collection comprises 89 Mat**a** strains and 46 Matα strains, as detailed in Table S1, a carefully chosen subset of the 1,011 isolates from Peter *et al*.’s [9]. Of these, 27 strains are present in our collection as both Mat**a** and Matα, resulting in a total of 108 unique strains. We begin with general characterizing of basic genomic and phenotypic properties our collection of strains. A substantial proportion of the strains (50) originated from the wine ecological niche, 12 strains are distributed across 8 different niches, each represented by 1 or 2 strains (**Figure 2A)**. Genomic analysis of these natural isolates resulted their phylogenetic classification into distinct clades. The majority of the strains (72) align with the wine clade or its sub-clades. Conversely, 14 strains are dispersed across 9 clades, each comprising only 1 or 2 strains **(Figure 2B)**.

**Figure 2.**
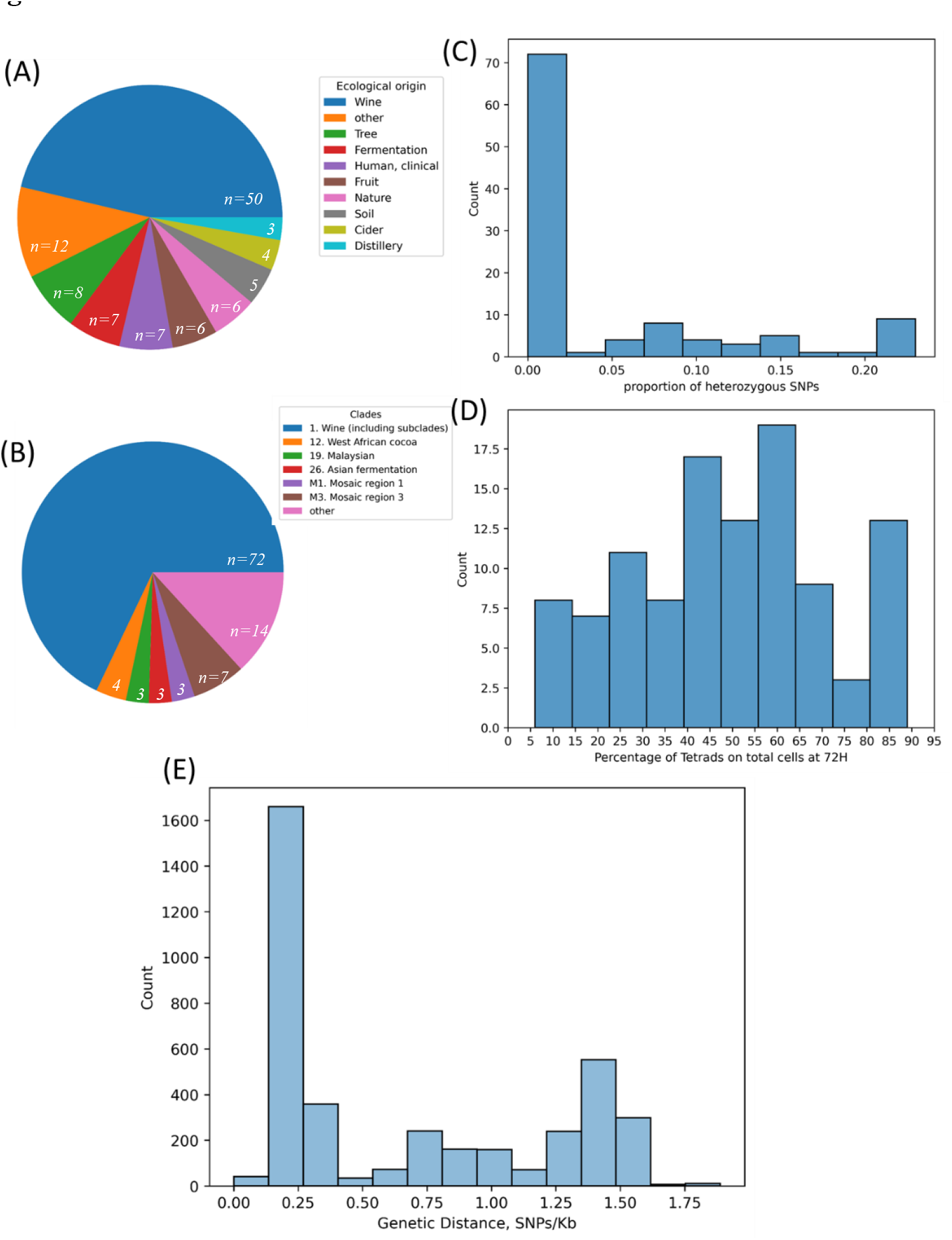
General characteristics of parental and h strains in this research. All data (apart from the sporulation data) is taken from Peter *et. al* 2018 [9]. sporulation data is taken from [38] (A-D) 108 unique strains were used in this analysis. (A) Distribution of ecological niches among the 108 parental strains. 8 ecological niches (Bakery, Beer, Industrial, Water, Flower, Human, Palm wine, Unknown) with 2 or less strains are shown together as “other”. Numbers in white color in the pie chart corresponds to the number of strains in this niche. (B) Distribution of clades among the 108 parental strains. 9 clades (7. Mosaic beer, 9. Mexican agave, 10. French Guiana human, 13. African palm wine, 18. Far East Asia, 21. Ecuadorean, 22. Far East Russian, 23. North American oak, 24. Asian islands) with 2 or less strains are shown as “other”. 2 strains have no clade assigned. All wine clades (including subclades) are shown together in the pie chart. Numbers in white color in the pie chart corresponds to the number of strains in this clade. Clade numbers are as in [9]. (C) Heterozygosity level distribution in the 108 parental strains (D) Distribution of sporulation efficiency as measured by number of tetrads seen after 72 hours of sporulation (E) Distribution of pairwise genetic distance. Pairwise GD was calculated for all crosses between 89 matA strains and 46 Matα strains

In the current study, we selected about 100 strains from the original collection 1,011 natural isolates. We employed several criteria in our strain selection. Firstly, we preferred strains that would be minimally heterozygote. The rationale of this criterion was that the more homozygote a strain is the higher is our confidence in the genetics of all its meiotic resultant haploids (See S**upplementary note 1** and **Figure 2C)**. Indeed, the majority of strains (70) exhibit a high degree of homozygosity with respect to most segregating SNPs, while the remaining strains display relatively low levels heterogeneity, with the maximum proportion of heterogenous SNPs being 0.35 out of the total SNPs segregating in the entire strain collection.

In addition, sporulation rate was originally measured for each strain as the proportion of tetrads after 72 hours of sporulation [39]. Sporulation efficiency in our collection vary between 5% and 95% **(Figure 2D**). Lastly, as part of the strain selection for this work we focused on a wide range of genetic distances (GD, as defined by the number of SNPs in 1Kb) (see also **supplementary note 1**). All pairwise GD were calculated for the 89 Mat**a** and 46 Matα and are plotted in a histogram in **Figure 2E**. Reassuringly, GD shows a multi-modal distribution with 3 discernable bins; between 0.05 and 0.5 SNPs/1kb, between 0.5 and 1.25 SNPs/1kb and above 1.25 SNPs/1kb. Additionally, we use a 0 SNPs/1kb bin as it represents a special case of Mat**a** and Matα originated from the same diploid natural isolate strain. Incidentally, the highest GDs between pairs of strains in this collection is about 1 SNP/kb which corresponds roughly to the SNP density between humans in the population.

### Fitness characteristics of parental and hybrids strains

We used the population of all hybrids and of all parents to measure fitness and study its modes of inheritance as a quantitative trait. The barcode fusion genetics systems (BFG) allowed us to mate all haploid parental strains *en masse*, potentially generating all hybrids combinations. In our quest to unravel the influence of environmental conditions on fitness inheritance, we conducted competition experiments using both a fermentable carbon source (glucose) and a non-fermentable carbon source (glycerol) which represents the two major energy metabolism modes in *S. cerevisiae*. Additionally, we introduced a non-metabolic stress condition, osmotic stress induced by sorbitol, in order to examine an abiotic stress in addition to the metabolic challenge. For the glucose and glycerol conditions, 3 repetitions were made; for glucose with the addition of osmotic stress 2 repetitions were made. Subsequently, we grew these hybrids in a competitive mode, enabling us to measure the relative fitness of each diploid descendant. We also subjected parents from each of the two mating types to competition, providing insights into their individual relative fitness.

Interestingly, not all hybrids pairs were generated (**Figure S3**). Across the three conditions 47% (∼2000) of possible hybrids were generated (an in depth analysis on mate pairing is described in Strauss *et al*. [40]). The fitness of each hybrid and of each haploid parent was determined by its frequency throughout the competition based on the logistic equation [37], [38]. Strains that became extinct during the competition (i.e., those observed at the beginning but eliminated later) or strains that were consistently absent throughout the entire competition were assigned low-fitness tags (see Materials and Methods).

Overall, the correlation between strain fitness in the repetitions of the same condition is ∼0.6. Clustering the conditions based on either the correlation between hybrid fitness, or simply the identity of hybrids that are made in each condition clearly separates between glycerol and the other two glucose-based conditions (**Fig S4**). Glucose and glucose with osmotic stress are clustered separately when looking on the identity of the strains made (**Fig S4B**) but not when considering the fitness correlation between the repeats (**Fig S4A)**. This suggests that in each of the three conditions, different hybrids were made (**Fig S4B**); but the same hybrids strains that were made in the different conditions perform similarly in the two glucose-based conditions (i.e. with or without sorbitol), and differently in the glycerol condition (**Fig S4A)**.

**Figure S5** shows a distribution of the relative fitness of all parents separately for each of the two mating types, and for all hybrids that had calculable fitness in each of the conditions (see Materials & Methods). We found a significant modest correlation between the relative fitness of the Mat**a** and Matα parents of the same strain on media containing glucose as carbon source (r=0.56; p-value ∼2*10^-2, **Figure S6**), indicating that, to an extent, fitness in these conditions is in part a property of the strain, shared between the two mating types. **Figure 3** shows the correlation between the mean relative fitness of the two parents of each strain and of the diploid hybrids, across the 3 growth conditions. The fitness of hybrids strains is mildly correlated between glucose and glucose with osmotic stress (r=0.38, p-value=2E-27) as well as between glucose and glycerol (r=0.27; p-value = 8E-27); glucose with osmotic stress is not correlated to glycerol (r=0.07, p-value=4E-2). Interestingly, we detect a significant correlation between mean parental fitness in glucose (with and without osmotic stress) and hybrid fitness in this condition and lack of such correlation in glycerol. Of note, we observe a correlation between mean parental fitness on glycerol and hybrid fitness on glucose and on glucose with sorbitol

**Figure 3.**
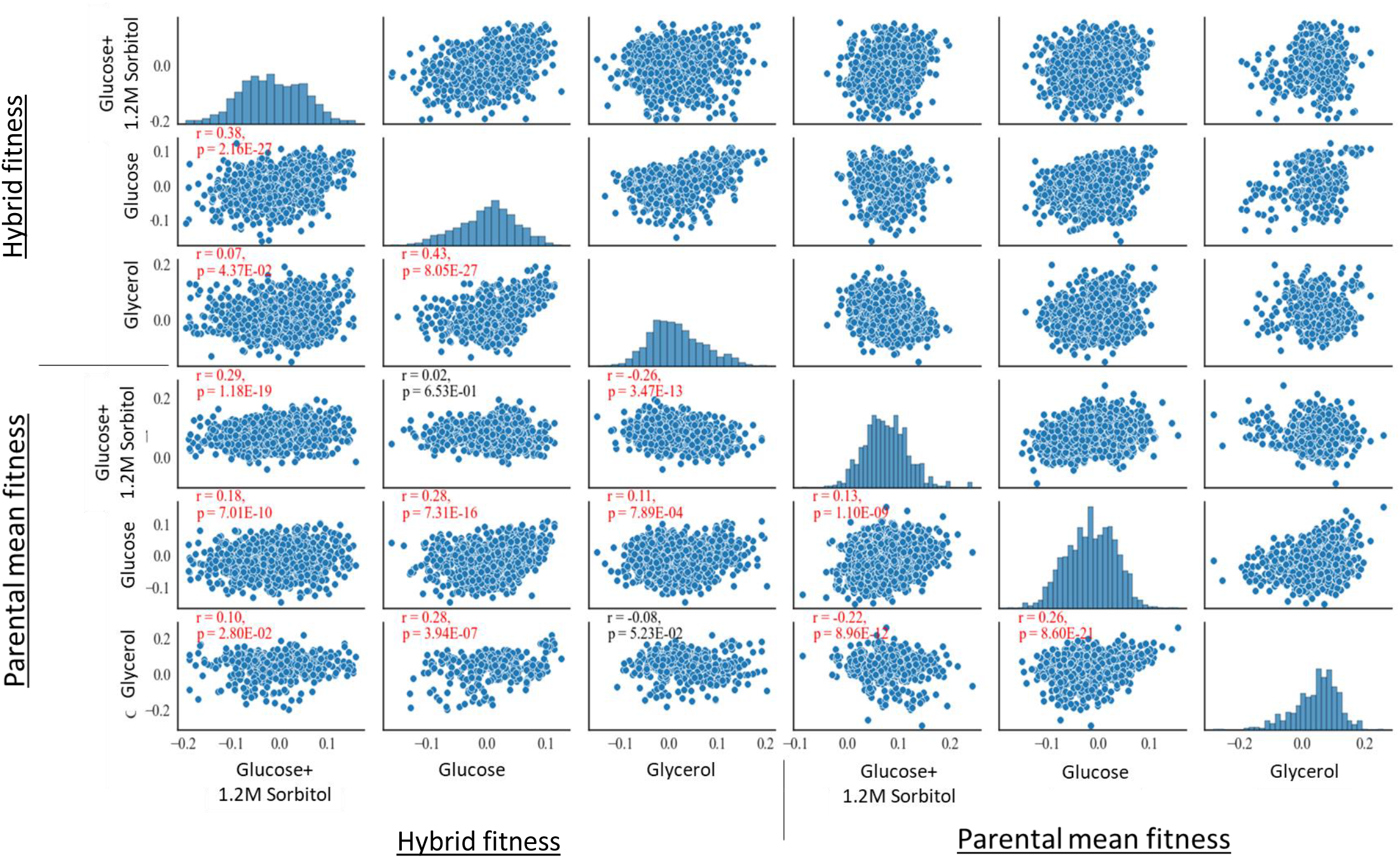
Distributions and correlations of mean parental fitness and hybrid fitness in the three conditions. Each sub plot represent comparison between two conditions. Dots represent either hybrid fitness or its mean parental fitness (top to bottom, left to right) in the three different conditions (Glucose+1.2M Sorbitol, Glucose, Glycerol, top to bottom, left to right). In the lower triangle, Pearson R as well as p-value is written on the scatter plot. Red color indicates a significant correlation. The diagonal shows the distribution of fitness values in the specific condition. The analysis was done on the continuous data only.

### Hybrid fitness correlates positively with parental fitness on glucose but less so on glycerol

We then wished to examine further if the fitness of hybrids correlates with the fitness of their parents under each growth condition. We begin by examining the correlation between hybrid fitness and the mean fitness of their parents. **Figure 4A** (and **Figure S7A**) shows a very different behavior between the fermentable and non-fermentable carbon sources. In glucose, parental fitness moderately but significantly correlates with hybrid fitness (r=0.27 p-value∼10^-15). In glycerol, there is no such correlation between the fitness of the parents and that of their hybrids. As an alternative to the mean parental fitness, we also characterized each pair of parents by their maximal or minimal fitness and computed the correlation between each of these parental measurements to hybrid fitness (**Figure S7B**,**C**). In agreement with the mean parental fitness, we find that in glucose hybrid fitness correlates positively and significantly with the maximal and with minimal parental fitness, but in glycerol such correlations were much lower. We also examined whether hybrid fitness correlated better with the fitness of the Mat**a** or Matα parent and found that the fitness of each of the two parental mating types correlated similarly with the hybrid fitness, and similarly to the correlation seen with the mean parental fitness, yet here too, only on glucose (**Figure S7D**,**E**). Thus, according to each of the parental fitness definitions, we find that in glucose (both with and without the osmotic stress) there exists a positive correlation with hybrid fitness while in glycerol we see little or no such correlation.

**Figure 4.**
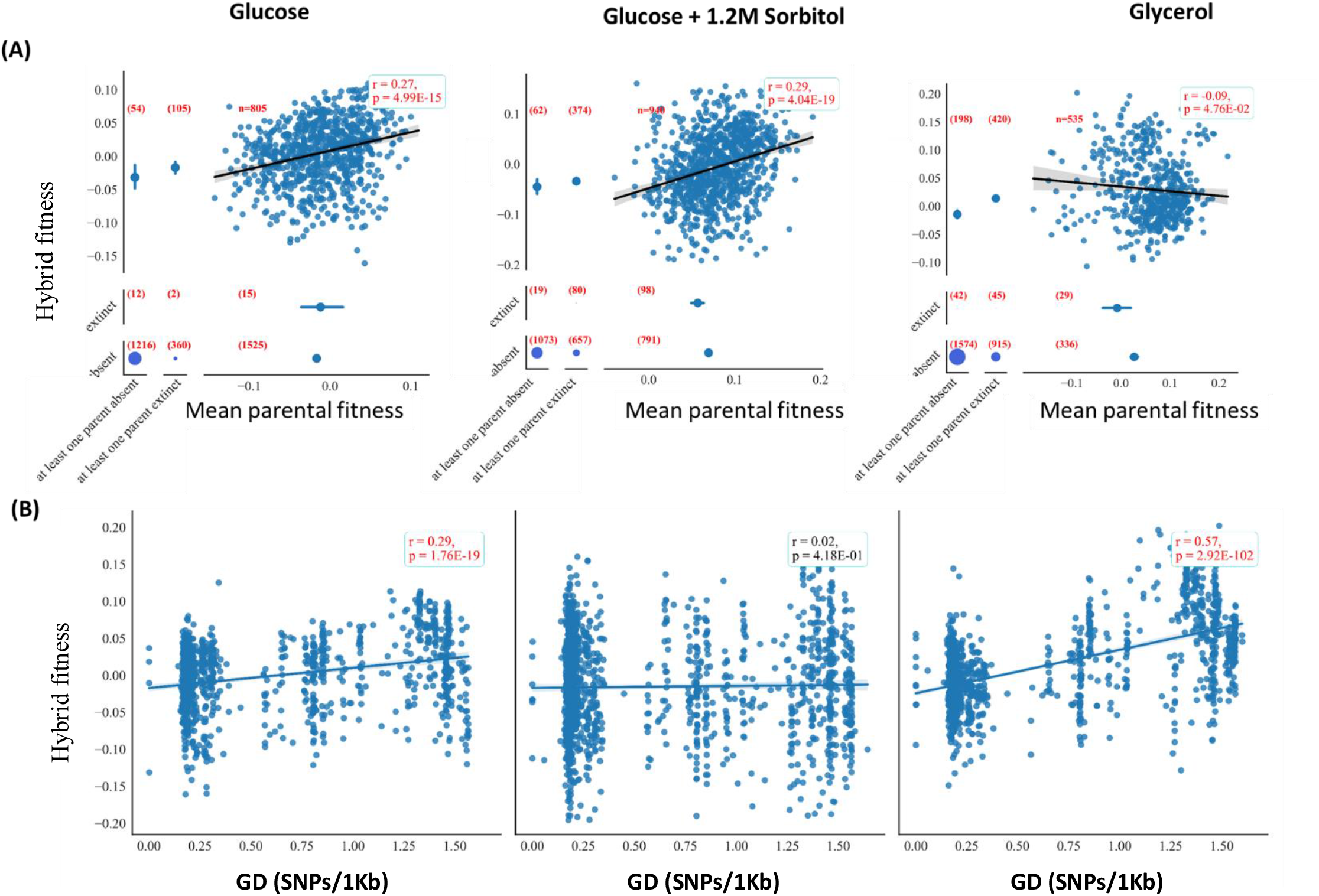
Fitness inheritance is mainly dependent upon parental GD in glycerol, but on parental mean fitness in glucose. Left: glucose as carbon source, middle: glucose as carbon source + osmotic stress (1.2M sorbitol), right: glycerol as carbon source (A) hybrids and parents’ fitness are positively correlated in glucose, but not in glycerol. In each plot the fitness of hybrids is plotted against the mean fitness of its two parents. Hybrids were partitioned according to their parents’ performance on the different carbon sources (X axis). Parent’s categories are (from left to right): at least one parent was absent in the competition, at least one parent was extinct during the competition, a continuous axis of mean parental fitness when the two parents survived. In addition, hybrids were partitioned based on their performance in the competition (y axis); hybrids either survived (top), were extinct (middle) or absent entirely (bottom). In the top right corner of each plot is a scatter plot and regression line of the calculable data only (hybrids that have survived and both of their parents has survived). Pearson correlation R and p value were calculated for each such plot and are shown in the box. The 4 circles in the bottom left of each plot are the number of hybrids found in each intersection of categories (i.e., hybrids that were either extinct or absent with at least one parent extinct or absent). Number of strains in each category is shown in red in brackets next to the relevant part of the figure. (B) Parental GD is positively correlated with hybrid fitness, mainly in glycerol. Each dot corresponds to an hybrid strain. Hybrid fitness (y axis) is plotted against the genetic distance between the two parents (x axis). The blue line represents the linear regression fit.

### An optimum, at high parental genetic distances, maximizes hybrid fitness, especially in glycerol

Each hybrid is defined by the genetic distance (GD) between its two parents. An interesting question is whether hybrid fitness is dependent upon parental GD. **Figure 4B** shows the correlation between hybrid fitness and their parental GD. In glycerol, hybrid fitness increases rather sharply with parental GD until it reaches a maximum at a distance of about 1.1 to 1.4 SNPs per kb. At higher GDs between parents, hybrid fitness starts to decline (**Figure S8**). The lowest hybrid fitness is observed among genetically close or identical parents, an observation that may be explained by homozygosity of recessive deleterious traits. In contrast, the high hybrid fitness at larger GDs may represent cases of beneficial heterozygosity in this condition. Yet, as parental genetic distance become even larger, the extent of heterozygosity may become maladaptive.

In glucose, the dependency of hybrid fitness on parental GD is much less pronounced (**Figure 4B**). Yet, in those conditions too (mainly without osmotic stress), hybrid fitness is maximized around a similar parental GD of about 1.2 to 1.4 SNPs per kb (**Figure S8**). Since yeast growth on non-fermentable carbon sources depends on mitochondrial function for aerobic respiration, we repeated all calculations of GD but with mitochondrial functional genes only, namely the set of genes encoded in the mitochondrial genome and mitochondrial genes encoded in the nuclear genome [41] and observed similar trends (**Figure S9A**). Namely, in glycerol parental GD, measured based on mitochondrial genes, correlates positively with hybrid fitness. We note that GD between parental pairs, calculated based on the entire genome, or based on mitochondrial nuclear genes only, are highly correlated (Pearson R=0.9, P-value ∼= 0) (**Figure S9B)**.

We have used a simple toy model of inheritance and found that a co-dominance mode of inheritance (where the heterozygot’s fitness is in between that of the two possible homozygots fitness) can explain the inheritance in glucose, while a dominance mode (where the heterozygot’s fitness is as high as the homozygot’s with the highr fitness) fits the inheritance mode in glycerol (see **Figure S10** and **supplementary note 2)**.

### High content of minor allele correlates with low fitness under a dominance genetic model, especially in glycerol

It is known that minor alleles (alleles that are rare in the population) are linked to low fitness (e.g., genetic disorders) in many genetic traits [42], [43]. As such, we asked if minor allele content of strains, summed over all SNP positions in the genome correlates with their fitness. To that end, we used the genetic annotation from Peter *et al*. [9] and estimated the content of minor allele homozygosity of each parental strain, and of each hybrid (see Materials & Methods). We used three models for this inference: (i) dominance, in which the heterozygous is as fit as the major allele homozygous, (ii) recessive, in which the heterozygous is as fit as the minor allele homozygous, and (iii) co-dominance in which the heterozygous has an intermediate fitness of the major allele and minor allele homozygous. For the dominance model only, there is a significant negative correlation between hybrid fitness and the parental minor allele contribution to fitness in glucose and glycerol but not for the glucose with osmotic stress condition. This correlation is stronger in glycerol (r=-0.26; p-value ∼10^-20 **Figure 5**). In contrast, in the co-dominance and recessive models, there is no correlation between hybrid fitness and parental minor allele contribution to fitness **(Figure 5**). In addition, by and large, no correlation is seen between parental haploid minor allele content and their fitness (**Figure S11**).

**Figure 5.**
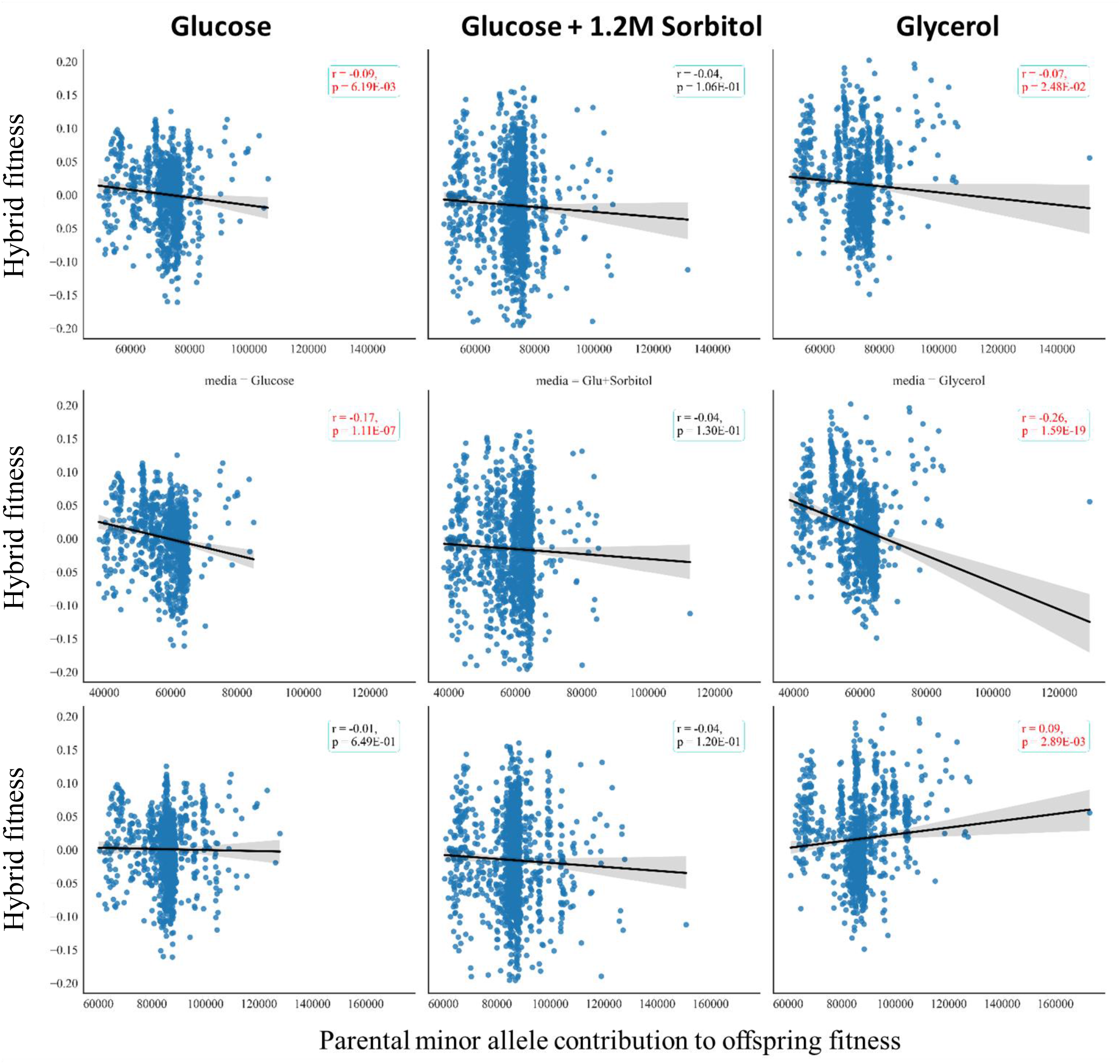
Hybrid fitness is negatively correlated to the contribution of parental minor allele content especially under the dominant model in glycerol. Hybrid fitness (y axis) is plotted against the contribution of parental minor allele content to hybrid (x axis, see Materials and Methods) weighted by the expected fitness under models of co-dominance (top), the dominance (middle) or the recessive (bottom) models. Each dot represents a hybrid and black line represents the linear fit. Experiments were performed on: glucose as carbon source (left), glucose as carbon source + 1.2M Sorbitol (middle) or glycerol (right).

### Hybrid fitness can be predicted using a multiple linear regression model

As shown so far, hybrid fitness is correlated to parental fitness, minor allele homozygosity content and parental GD (although to different extents in the two carbon sources). Therefore, we next asked if hybrid fitness can be predicted based on those three parameters using a linear model (multiple linear regression, see Materials and Methods). We compared the correlation between the observed hybrid fitness to the fitness of the hybrids as predicted by our model (**Figure 6)**. We succeeded to predict hybrid fitness with high accuracy (as depicted by the correlation to the measured fitness) in glycerol and moderately yet significantly in glucose (r=0.71 and r=0.38 respectively in the two conditions). In agreement with the above findings, the model coefficients show that GD is mostly important in predicting fitness in glycerol, while the homozygosity minor allele content and parental mean fitness contributes more to the prediction in glucose as carbon source. These results indicate that in glycerol about 50% of the variance in hybrid fitness is explained by parental GD.

**Figure 6.**
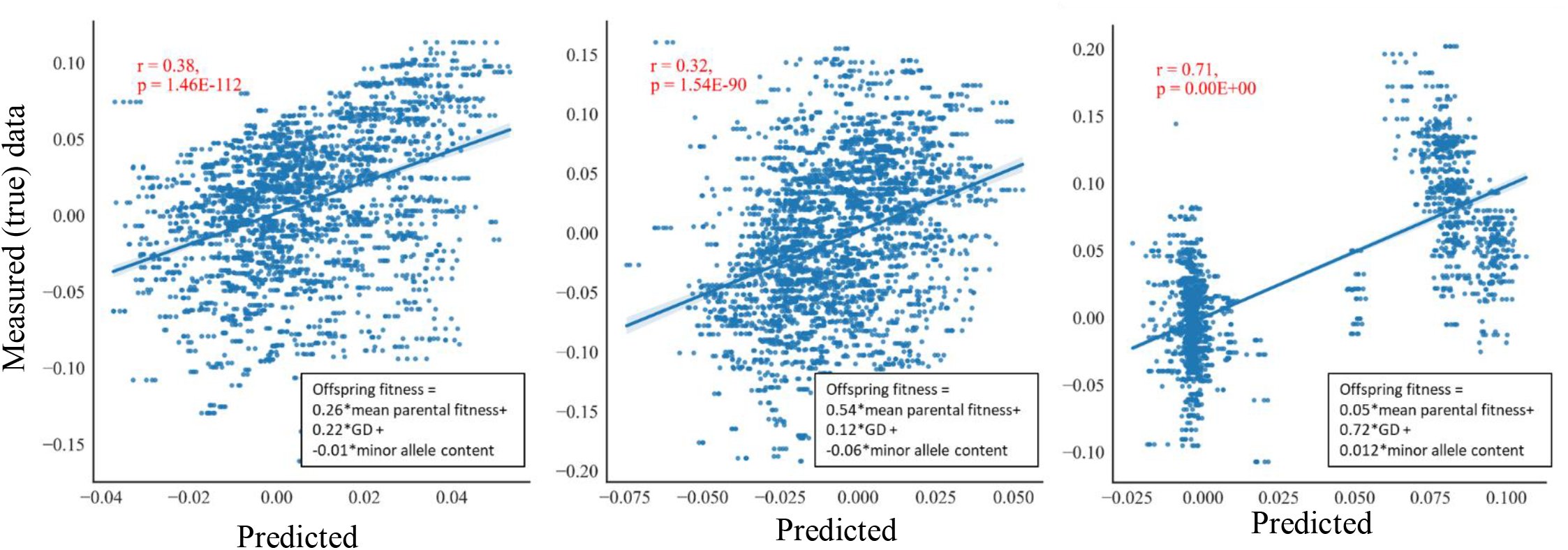
Multiple linear regression model predicts hybrid fitness based on parental fitness and parental GD. Left: glucose as carbon source, middle: glucose as carbon source + osmotic stress (1.2M sorbitol), right: glycerol as carbon source Multiple linear regression (MLR) estimation for hybrid fitness based on parents’ mean fitness, minor allele content and parental GD were performed. A multiple linear regression model was calculated based on a training set containing 80% of the data, the plots shown here apply to 20% that served as a test data. The procedure was repeated 20 times, and the test set results were aggregated to make the figure. Model parameters are shown in the equation in the bottom right hand corner (coefficients were averaged over the 20 repeats of MLR). Each point represents a strain (from the test data). X axis is the expected fitness value by the model, while the y axis represents the experimentally measured fitness value of the strain. Pearson correlation and p-value between the prediction and true value are shown in red in upper left corner. Analysis in this figure was done on continuous data only (hybrids and parents that were not extinct during the experiment). Predicted fitness in glycerol appears to be bi-modal, probably reflecting the fact that GD distribution is mainly bi-modal and that this variable is the main contributor to the predicted fitness in this condition.

## Discussion

We present here a new resource, genomic methodology, sharable biological material and a large database, for the tracking of mating and fitness measurement of sexual mating hybrids in yeast. We have harnessed the inherent genetic diversity and fitness within the natural *S. cerevisiae* isolates collection [9] to establish a platform for investigating questions related to genetics and inheritance of fitness as a quantitative trait. By implementing a barcoding system that fuses barcodes in the diploid hybrids resulting from the mating of two natural isolates each with its own barcode, we can generate thousands of variants while simultaneously monitoring mating pairs.

In this study, we utilized this platform to explore modes of inheritance between parents and their hybrids of fitness as a quantitative trait. Our findings reveal two distinct modes of inheritance. In fermentable carbon sources, such as glucose, hybrid fitness correlates with parental fitness, with a lesser dependence on the genetic distance between them. Conversely, in a respiratory condition, with glycerol as a carbon source, hybrid fitness is predominantly influenced by the genetic distance between the parents, and it is typically higher with higher genetic distance. We have chosen 108 (89 Mat**a** and 46 Matα) parental strains from the 1,011 strains in the entire strain collection, as those that are minimally heterozygote (see [9], **Figure 2** and **supplementary note 1)**. When perfectly homozygote, parents can inherit only one specific set of chromosomes. When this holds, parental genetic distance translates uniquely to a certain level of heterozygosity among the hybrids – any SNP between the two homozygote parents will lead to one site of heterozygosity in their hybrids. Since our parental strains were largely homozygotes, we can deduce that hybrids whose parents were separated by a large genetic distance are more heterozygote on average than hybrids whose parents are close. Thus, the strong positive correlation between fitness on glycerol and parental genetic distance, also suggests that high heterozygosity of hybrids is correlated with high fitness at that condition.

A simple genetic model may explain the broad range of observations we make by assuming one of several possible modes of inheritance: in the co-dominance model heterozygous fitness falls between the fitness of the two homozygous types, and the dominance model, where heterozygotes exhibit fitness equivalent to the better homozygous type. Our results are in agreement with the predictions of the dominance model in glycerol and the co-dominance model in glucose. Further investigation will be needed in order to explain these trends. Since fermentation is the favored condition for most strains, the non-fermentative glycerol carbon source might be seen as a type of stress. Yet, interestingly, when we grew the cells on glucose as fermentative carbon source, but applied an osmotic stress, the modes of inheritance of fitness resembles those in glucose and not in glycerol. This suggest that not stress *per se* determines modes of inheritance, but rather the major dichotomy between fermentation and respiration.

Our system provides a general resource for the study and application of yeast quantitative genetics. While our focus here was on assessing fitness inheritance, this platform holds broad applicability for investigating the inheritance of various other traits such as gene expression level, and other cellular aspects of yeast cells such as size. For instance, by measuring intensity of the fluorescent proteins (GFP and RFP) in parents and in hybrids, we can unveil trans effects on gene expression in yeast (manuscript in preparation). Given the platform’s capability to create and track numerous isolates, it opens avenues for addressing other questions, such as mate choice. By investigating the pairing established between parents and their frequency in the population, we can address inquiries related to mate choice and its impact on fitness (Strauss *et al*. [40])

## Materials and Methods

### Strains and media

Strains in this project are natural isolates taken from Peter *et al*. [9]. **Table S1** shows the list of strains used in this work.

The following media types were used:

**SD Glu–** 6.7g/L yeast nitrogen base, 1.5g/L amino acid mix and 2% Glucose (according to [72]).

**SD Gly–** 6.7g/L yeast nitrogen base, 1.5g/L amino acid mix and 2% Glycerol (according to [72]).

**SD Glu + 1.2M Sorbitol** -6.7g/L yeast nitrogen base, 1.5g/L amino acid mix, 2% Glucose and 1.2M Sorbitol

**YPD -** 10g/L yeast extract, 20g/L peptone, 20g/L glucose

**YPA** - 10g/L yeast extract, 20g/L peptone, 20g/L Potassium Acetate

**SPO media -** 2.5g/L yeast extract, 15 g/L potassium acetate Antibiotic concentrations and initials:

Hygromycine B (Hyg) - 300mg/L,

Nourseothricin (NAT) – 100mg/L,

Kanamycin (G418) – 200mg/L

Zeocin (Zeo) – 200 mg/L

### Plasmids and constructs design and synthesis

Strains were engineered by integrating into their genome a compound design that included the following parts; (**Figure S2**) (i) genome homology region: 200bp and 700bp homology to the HO locus on both ends of the construct. The specific sequences of homology were chosen to have the least amount of SNPs between most strains, using multiple sequence alignment and BLAST search on strains’ genomes published in Peter *et al*. [9]. Homology region was amplified from the genome of BY4741 strain (upstream homology region, chrIV:46062…46271, downstream homology region, chrIV: 47982…48682, (ii) Barcode Fusion Genetic (BFG) system (**Figure S2B**). This region includes (1) the barcodes region flanked with lox sites that was synthesized and cloned into the rest of the construct by outsourcing company (Twist bioscience). The barcode region was designed based on [33]. (2) the Cre enzyme and rtTA inducer (**Figure S2A**). This region was kindly given to us by the lab of Fredrick Roth [33], (iii) Constitutive markers and fluorescence proteins (kindly given to us by Naama Barkai’s lab). Two combinations of constitutive fluorescence and antibiotic resistance was used, either a yeGFP gene under the control of the TDH3 promoter and Hygromycine resistance cassette with a TEF1 promoter (in Mat**a** construct), or a mCherry gene under the control of TEF2 promoter and Nourseothricin resistance cassette with a TEF1 promoter (in Matα construct). (iv) Haploid selecting markers. Zeocin or Geneticin (G418) resistance cassettes under the control of the Ste2 Mat**a** specific promoter or Ste3 Matα specific promoter, respectively. A synthetic terminator was added to both resistance cassettes [44]. Sections (iv) was synthetized by outsourcing company (GenScript).

All of the above parts, except for (ii) were assembled together (in the order described in **Figure S2**) using restriction free methods and cloned into pET28a (Novagen #69864-3) plasmid by the cloning unit in Weizmann institute, generating one backbone plasmid to generate Mat**a** strains and another for Matα. The barcode fusion genetics library section (ii1) was then cloned into the Mat**a** and Matα plasmids by Twist Bioscience Company. **Figure S2A**, top panel shows the construct that was transformed to create Mat**a** cells, while the bottom panel shows the construct that was transformed to create Matα cells (from now on referred to as Mat**a** construct and Matα construct, respectively).

### BFG construct and primers

The BFG system (shown in **Figure S2B)** is composed of barcodes flanked by *lox* sites (based on [33]). While in the Mat**a** construct the order is *loxP-BC1-lox2272-BC2*, in Matα construct the order is opposite *BC1-loxP-BC2-lox2272*. The other main component is the Tet-on system that is composed of an rtTA inducer under constitutive promoter and a *Cre* enzyme under pTet promoter. The rtTA inducer is only active, and can mediate Cre transcription when tetracycline presents.

Following activation of Cre enzyme, a recombination event takes place, recombining the barcode regions of the two parents and results in two fused fragments, one on each chromosome; *BC1(Mat****a****)-loxP-BC1(Matα)-lox2272* and *loxP-BC2(Mat****a****)-lox2272-BC2(Matα)*

In addition to the barcodes and *lox* sites, these regions contain unique sequences of ∼25nt that are either shared between the parents or are unique to each construct. Those regions can thus be used as primers for amplifying either Mat**a** barcodes construct only (**primers H and K**), Matα barcodes construct only (**primers B and F**), diploid hybrids fused barcodes only (**primers B and I** or **primers E and K**), or all of the above (**primers A and G**).

Primers used for amplifying Mat**a** construct only, Matα construct only, fused barcode only or all, will be termed Mat**a** primers, Matα primers, fused primers or general primers, respectively. Primers sequence can be found in **Table 1**.

**Table 1.**
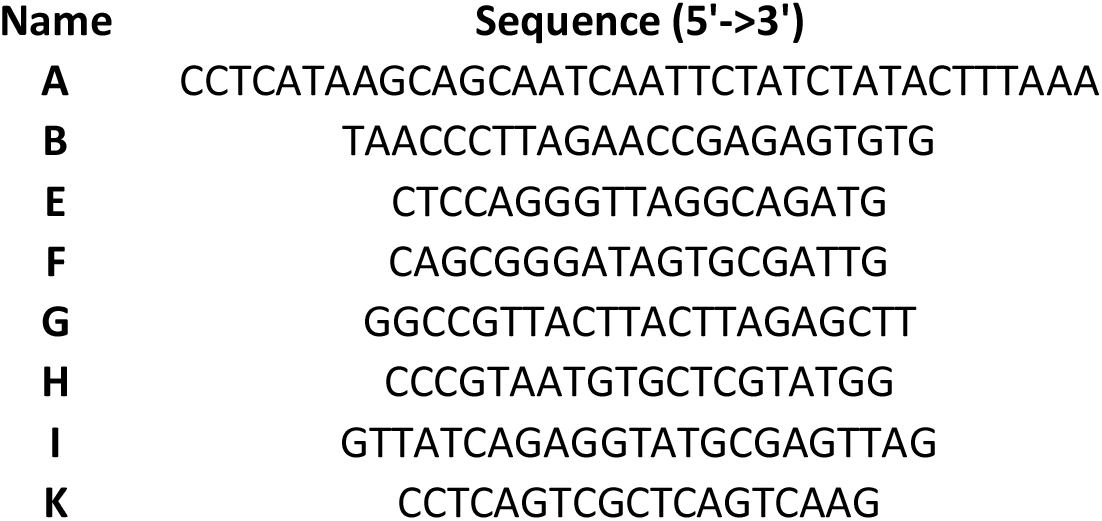
List of primers of the BFG system used in this paper

### Strain construction

#### Transformation

Constructs (from the section *“Plasmids and constructs design and synthesis”*) were amplified using the following primers; F: GGTGAAAACCTGTACTTCCAGGG, R: ATGCTAGTTATTGCTCAGCGGT. PCR was done using KAPA HiFi HotStart ReadyMix (Roche, KK2602) enzyme according to manufacturer instructions with the following details: primer annealing Tm of 64°C, elongation of 5 minutes, 30 cycles. PCR products were then transformed into the chosen strains as follows: For each transformation, 5 reactions of PCR were made (to increase complexity of the library). All reactions were ran on agarose gel (0.8% agarose) and size was verified (7kb).

To transform the cells, LiAc protocol [45] was adapted to allow high throughput transformation of many strains in a 96-well plate. In short, 48 strains were taken out of the -80 and inoculated in a 96-well plate in a checkerboard manner to avoid cross contamination between strains. Cells were grown overnight at 30°C with shaking (1200rpm), in a shaker incubator. Then, each strain was diluted 1:20000 in a 50ml tube, (0.5ul of culture into 10ml YPD) and grown for 16 hours at 30°C while shaking. Following the growth phase, four random strains were counted to estimate cell concentration. Cultures usually reached the late log stage (budded yeast, ∼7E7cells/ml); each culture was diluted 1:50 into fresh YPD and allowed to grow for another couple of hours.

Cells were harvested by centrifugation (4000rpm for 5min) and washed twice; first with 5ml DDW then with 1ml LiAc 100uM. After the second wash, remaining liquid was vacuumed, and pellet was re-suspended in 60ul DDW. Two 96-well plates for transformation, in a checkerboard manner (one for Mat**a** construct and one for Matα construct) were made with 25ul of PCR product of the relevant construct. 30ul of each strain were suspended into each of the two transformation plates. Transformation mix (100ul PEG 50%, 15ul LiAc 1M, 4ul of 10mg/ml salmon sperm (Sigma Aldrich, D9156-1ML) were added to each well. Plates were incubated for 40 minutes at 42°C. Following incubation, plates were centrifuges (3000rpm for 3 minutes), and liquid was vacuumed with a multi pipette vacuum adaptor. 150ul of YPD was added to each well and plates were incubated overnight at 30°C. The following morning, each well was plated on YPD agar containing the relevant antibiotic (Hyg for Mat**a** construct plate, and Nat for Matα construct plate). Agar plates were incubated in 30°C for a couple of days until the appearance of colonies.

After colonies appeared, four colonies from each strain were picked, inoculated into liquid SD+Glu in 96-well plate (to verify fluorescence) and patched on YPD agar plate with the corresponding antibiotic (for continuing). 96-well plates were grown overnight at 30°C and diluted 1:50 into fresh SD+Glu. Plates were FACS analyzed to verify the correct fluorescence marker (cells with Mat**a** construct had yeGFP while Matα construct corresponds to mCherry). One positive colony per strain was chosen to continue.

### Sporulation and random spore analysis

To generate engineered strains, positive diploid colonies (obtained as described in the section “transformation”) were inoculated into a 24-well plate with 1.2ml YPA and grown overnight at 30°C with shaking (1200rpm). Plates were centrifuged (4000rpm, 3 minutes) and washed with DDW, spin down and vacuumed. Pellets were re-suspended in 1.2ml SPO media and incubated for 4-5 days in 25°C while shaking. After 4 days, a couple of random cultures were observed under the microscope to verify sporulation, and score sporulation efficiency. In case of low sporulation efficiency plates were incubated an extra day.

50ul of each sporulated culture was transferred into a 96-well plate for random spore analysis as follows:

Cultures were centrifuge and pellets were re-suspended with 50ul TE buffer (Tris 10mM EDTA 1mM, pH 8.0) supplemented with 2.5ul β-Mercaptoethanol (final concentration of 0.1M) and incubated for 10 minutes at room temperature. Plates were centrifuged and liquid was discarded, pellets were then washed with 150ul DDW twice. Pellets were re-suspended in 50ul β-Glucoronidase (Sigmacat no. G7017-5ML dilute 1:2 and filtered) and incubated for two hours in 37°C, while shaking. Plates were centrifuged and pellets were washed with 200ul Triton-X100 0.5%, this step was repeated twice. Plates were centrifuged, and pellets were re-suspended with 120ul DDW and plated on YPD agar plates, with corresponding antibiotic for haploid of the correct mating type (Mat**a** construct were plated on Zeocin containing plates, while Matα construct cells were plated on G418). Plates were incubated in 30°C for 48h until colonies appeared.

From each strain, 2 colonies were picked, inoculated into liquid SD+Glu in 96-well plate and patched on YPD agar plate with the corresponding antibiotic. 96-well plates were grown overnight at 30°C and diluted 1:50 into fresh SD+Glu. Plates were FACS analyzed to verify correct fluorescence marker (cells with Mat**a** construct had GFP while Matα construct corresponds to mCherry).

Positive colonies (1-2 colonies) were continued to Sanger sequencing to recover barcode sequences.

One correct haploid colony per strain was frozen in a 96-well plate in -80°C.

### En masse mating

All verified haploid strains were taken out from the -80°C into YPD media with corresponding antibiotic (Hyg or NAT for Mat**a** or Matα respectively) in a 96-well plate using pinners. Strains were grown overnight (30°C, while shaking) and then diluted 1:1000 for another overnight incubation in 30°C. Strains were diluted 1:50 into either SD-Glu or SD-Gly and grown for another couple of hours in 30°C to reach mid-log phase. OD was measured using plate reader (infinite 200, Tecan). All Mat**a** strains and all Matα strains were mixed (separately for Mat**a** and Matα), based on the measured OD, such that they will have equal representation in the mix. Mixes were centrifuged and re-suspended in 0.2X volume to create a 5-fold increase in cell concentration (∼1E8 cells/ml). *En masse* mating was conducted by mixing 50ul (∼5E6 cells total) of each mating type mix in a 1ml medium (either SD-Glu or SD-Gly) with doxycycline (final concentration of 10ug/ml). Mating was executed for 20 hours in 25°C without shaking.

### Pooled competition

For measuring fitness of each hybrid we conducted pooled growth competitions. For that, each of the *en masse* mating was continued to a competition assay. Competitions were done in a daily dilution (1:240) manner in either SD-Glu, SD-Glu + 1.2M sorbitol or SD-Gly, with both Hyg and NAT to select for diploids. In the first dilution, media also contained doxycycline (10ug/ml). Cultures were grown in 50ml tubes at 10ml volume in 30°C while shaking. Dilution was carried out every day (in SD-Glu) or every two days (SD-Glu + 1.2M sorbitol and SD-Gly) by transferring ∼40ul of the culture into 10ml of fresh media.

Cells were frozen in 30% Glycerol and kept in -80°c every dilution (8 generations per dilution, for a total of ∼55 generations). All but the first dilutions were freezed leading to 6 time points per a competition experiment.

To determine fitness of the parents, competition experiments for all parents of a specific mating type (either Mat**a** or Matα) were conducted similarly (separately for each mating type).

Frozen samples were used for library preparation, sequencing and fitness calculation using the following time points; hybrids (diploids) – generations 16, 24, 32, 40, 48, 56; haploids (parents): generations 8,16,24,32,40.

Parental competitions in all 3 media types as well as the hybrids competition on Glu+1.2M Sorbitol were done in two repeats; hybrids competition on SD-Glu and SD-Gly were done in three repeats each.

### Library preparation, sequencing and read analysis

DNA was extracted using MasterPure Yeast DNA Purification Kit (Epicenter), according to the manufacture protocol.

Libraries for sequencing the barcode region were constructed by designing Plate-Row-Column PCR methodology; in which a first PCR is done using primers targeting the barcode region as well as adding plate barcode and tails that match Illumina adapters (F: ACGACGCTCTTCCGATCT**NNNNN***BFGprimer*, R: AGACGTGTGCTCTTCCGATCT**NNNNN***BFGprimer*)

Capital letters corresponds to Illumina adaptors, N correspond to plate index and BFGprimer is the primer shown in Table 1. First PCR was done in 25ul final volume with 2ul of template DNA (∼100ng genomic DNA). PCR program: Tm of 60°C, elongation of 10 seconds, ∼20 cycles. Usually, 4 PCR reactions were done per experiments and they were pooled together after the first PCR (to avoid PCR biases). 2ul of the first PCR was used as a template for the second PCR.

A second PCR with the following primers (F: AATGATACGGCGACCACCGAGATCTACACTCTTTCCCTACACGACGCTCTTCCGATCT, R:CAAGCAGAAGACGGCATACGAGAT***NNNNNNNN***GTGACTGGAGTTCAGACGTGTGCTCTTCCGATCT. ***N*** corresponds to Illumina index for library multiplexing) was carried out to attach the adapters for the Illumina run. PCR was done in 25ul volume. PCR program: Tm of 62°C, elongation of 10 seconds, ∼20 cycles. After second PCR libraries were cleaned using SPRI beads (∼1.2X ratio) to eliminate unspecific bands and primer dimers.

Amplicons were sequenced using paired-end methodology, on the NovaSeq platform (Illumina) After initial de-multiplexing by the Illumina platform, libraries sequencing reads were further separated based on the plate index using cutadapt [46]. All reads were further processed by cutadapt to leave only the barcode region. For alignment, a synthetic genome from all strains’ barcodes was made using bowtie2 (bowtie2-build command). Alignment was performed using bowtie2 as well. Read counts per variant were determined by in-house script.

### Fitness estimation based on pooled competition

Fitness was derived by employing a Maximum-Likelihood (ML) algorithm on all read count measurements along the competition experiment per variant. Briefly, first, each variant fitness is estimated by using a simple loglinear regression over the first three time points. Based on these estimations, the initial relative frequencies of each variant, and a noise model that accounts for experimental errors [37], [38], expected trajectory of each variant is estimated and compared to the measured trajectory. Next, small changes are made to fitness estimates, comparison is repeated, fitness is updated if they better fit the data (higher likelihood). This procedure is performed iteratively until fitness estimates are stable (maximized likelihood).

We measured relative fitness as vegetative growth rate of each of the haploids from each mating type at the presence of all others, and of each of the diploid hybrids at the presence of all others. Measurement of the relative fitness of each strain in each such pool is performed by deep sequencing of the parental (for haploids) and recombined (for diploids) barcodes in pooled competitions, in similarity to previous studies [37], [38], [47]. Relative fitness was measured and calculated separately for each condition. Fitness is derived from the following equation:

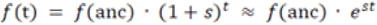

The frequency of a strain at time point *t* is a product of its initial frequency and its relative fitness, raised to a time-dependent power (f = frequency of a strain at a given time point, t = time, anc = strain in time point zero, s = fitness coefficient). Overall, we performed the competitions over 40 and 56 generations for the haploids and diploids respectively.

### Filtering strains and re-assigned fitness

After initial fitness calculation, strains were filtered, and for some strains fitness was re-calculated (according to the criteria describes below). Fitness distribution was normally distributed (between -0.2 and 0.2) with an additional single peak (usually at -0.6 or -0.8 that was detected as variants that were extinct during the competition).

#### Hybrid (diploid) fitness re-assignment

First, a goodness of fit (GoF) was calculated per variant as follows; PyFitSeq algorithm outputs the calculated read counts in each time point, given the initial read counts in the first time point and the calculated fitness. For each variant, GoF was determined as the Pearson coefficient between the calculated read counts and the real read counts data. Strains were filtered to have GoF>0.5 (∼75% of the data in SD-Glu, ∼90%SD-Gly, 65% of the data in SD-Glu + 1.2M sorbitol). In addition, variants were filtered to have at least 100 read counts in the first time point, or having more than 10 read counts in 3 additional time points (∼50% of the data). Next, a fitness per variant per media was calculated as follows; (i) if the variant was not extinct in any of the repetitions per media, fitness of variant was determined to be the average of the repeats (ii) if variant was extinct in more than 60% of the repetitions per media, variant was determined to be “extinct” (arbitrary fitness of -1), (iii) if variant was extinct in less than 40% of the repetitions per media, the extinct value was removed, and fitness was determined to be the average fitness in the other repetitions (iv) if variant was extinct in 40% to 60% of the repetitions fitness was calculated for each variant in each repetition. In case this score was <-0.4, fitness value in that repetition was changed to “extinct”, then, if the variant was extinct in >60%, variant was determined to be “extinct” (arbitrary fitness of -1), (v) lastly, a few variants could still not be given a single value. Those variants were inspected manually, showed a genuine decrease in read counts even if not extinct, so they were determined as “extinct”, but given an arbitrary fitness value of -2.

#### Parents (haploid) fitness re-assignment

First, strains were filtered to have at least 200 reads in the first time point (∼80-90% of data). For filtered variants, fitness was calculated similarly to the hybrids calculation; (i) if the variant was not extinct in any of the repetitions per media, fitness of variant was determined to be the average of the repeats (ii) if variant was extinct in more than 50% of the repetitions per media, variant was determined to be “extinct” (arbitrary fitness of -1), (iii) if variant was extinct in 25% of the repetitions per media, the extinct value was removed, and fitness was determined to be the average fitness in the other repetitions, (iv) lastly, a few variants could still not be given a single value. Those variants were inspected manually, showed a genuine decrease in read counts even if not extinct, so they were determined as “extinct”, and given an arbitrary fitness value of -1. One strain (Matα, AIM in SD-Gly) showed no read counts in all time points (except for the first one) and it was determined as “absent”.

### Homozygous minor allele content calculation

Relevant strains were selected from the .gvcf file [9]. For each parental strain, +1 was added to the minor allele content if original strain was homozygous for minor allele, or +0.5 was added if original strain was heterozygous. For hybrids, three models were used to predict hybrid fitness based on parental minor allele content (**Figure 6**) (also see **Table 3**); (i) in the dominance model, +1 was added to the minor allele content if the two original parental strains were homozygous for minor allele, +0.5 was added if one strain was heterozygous and the other was homozygous for minor allele, and +0.25 was added in both parental strains were heterozygous for the minor allele. (ii) co-dominance model: +1, +0.75 and +0.5 were added to the minor allele content respectively, (iii) recessive: +1, +1, +0.75 were added to the minor allele content respectively. Those numbers take into consideration both the chances of the hybrids to be either homozygous or heterozygous for the minor allele, as well as the additive value of the genotype.

**Table 3.** Calculation of parental minor allele content to hybrid fitness under different genetic models

| Parental genotype<br>(vcf notation, genetic notation) | Chance for homozygous minor allele | Chance for heterozygous | Dominance (homozygous *1) | Co-dominance (Homozygous*1 + heterozygous*0.5) | Recessive (Homozygous * 1 + heterozygous * 1) |
| --- | --- | --- | --- | --- | --- |
| (0,0)/(0,0) AAXAA | 0 | 0 | 0 | 0+0 | 0 |
| (1,1)/(1,1) aaXaa | 1 | 0 | 1 | 1+0 | 1 |
| (1,1)/(1,0)\|(0,1)aaXAa | 0.5 | 0.5 | 0.5 | 0.5+0.25 | 0.5+0.5 |
| (1,0)\|(0,1)/(1,1)AaXaa | 0.5 | 0.5 | 0.5 | 0.5+0.25 | 0.5+0.5 |
| (1,0)\|(0,1)/(1,0)\|(0,1) AaXAa | 0.25 | 0.5 | 0.25 | 0.25+0.25 | 0.25+0.5 |

### Calculating mitochondrial genome-based parental GD

Gene list of 250 mito-nuclear genes were downloaded from the *Saccharomyces* genome database website (SGD) [48]. Eight additional mitochondrial genes were found in the 1,011 strains website (http://1002genomes.u-strasbg.fr/files/). For each pair of strains, pairwise sequence alignment was performed using python Bio package (Bio.pairwise2.globalxx). For each gene, for each pair of strains #SNPs/total length of gene was calculated and saved. Final mitochondrial GD was calculated as the sum of all SNPs frequencies from all genes, for each pair of strains. all genes sequences were obtained from *Peter et al*. [9].

## Acknowledgements

We thank the Pilpel lab for extensive feedback and discussions. We thank Prof. Naama Barkai for useful comments. We thank Prof. Frederick Roth from the University of Toronto for sharing their BFG design and thoughts with us.

All cloning, but adding the BFG into the construct, was done by Dr. Yoav Peleg from the Weizmann Institute core facilities.

Sequencing was performed at the Crown Institute of Genomics, G-INCPM at Weizmann Institute of Science.

We thank the Israel Science Foundation, the Yeda-Sela Foundation at the Weizmann Institute, and the Minerva Foundation for grant support. YP is the Ben-May Professorial Chair and a Kimmel Investigator.

