## Supplementary Figures for "Quantitative genetics of natural *S. cerevisiae* strains upon sexual mating reveals heritable determinants of cellular fitness"

### Slide 1
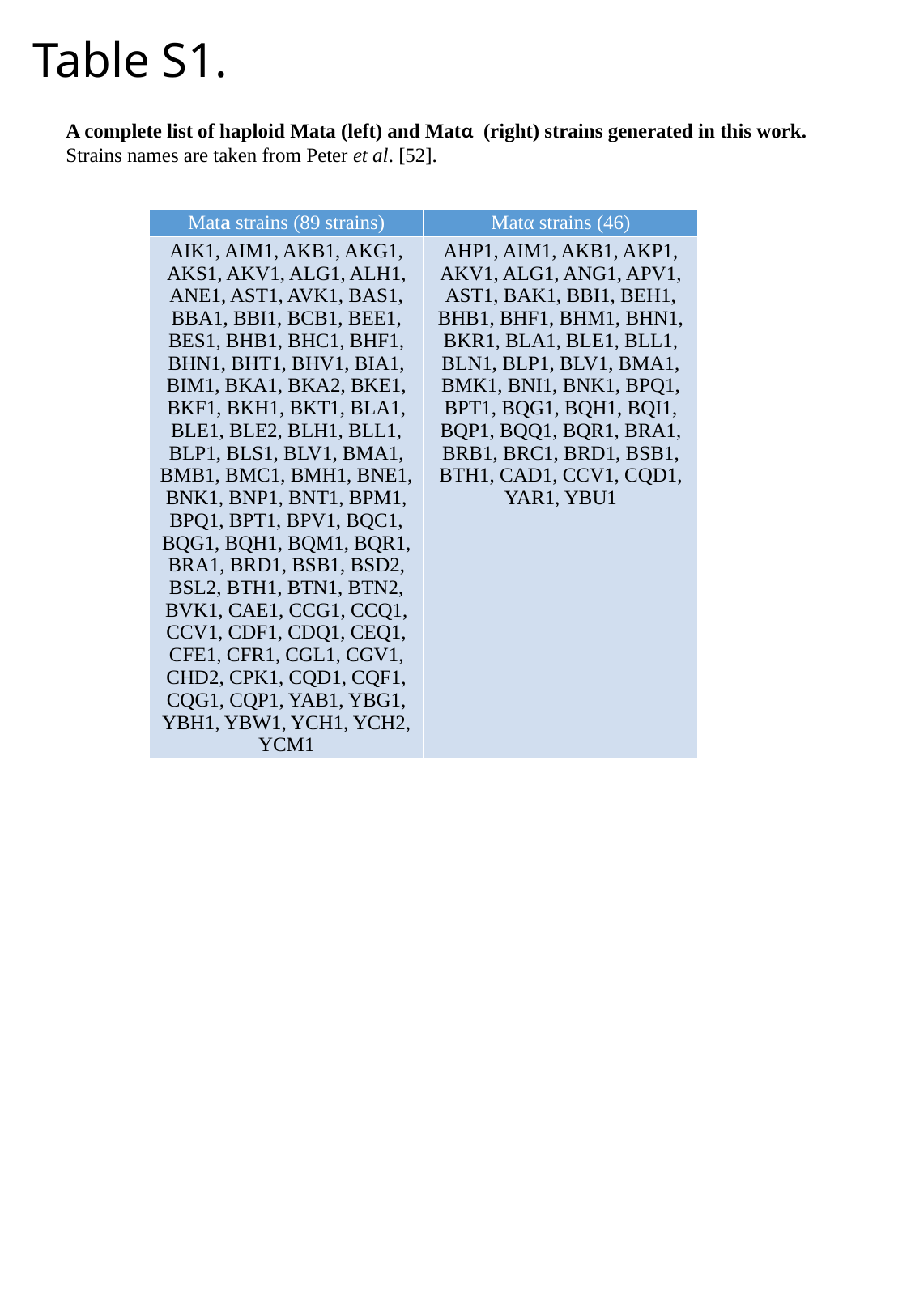

#### Table S1.
A complete list of haploid Mata (left) and Matα  (right) strains generated in this work. Strains names are taken from Peter et al. [52].
| Mata strains (89 strains) | Matα strains (46) |
| --- | --- |
| AIK1, AIM1, AKB1, AKG1, AKS1, AKV1, ALG1, ALH1, ANE1, AST1, AVK1, BAS1, BBA1, BBI1, BCB1, BEE1, BES1, BHB1, BHC1, BHF1, BHN1, BHT1, BHV1, BIA1, BIM1, BKA1, BKA2, BKE1, BKF1, BKH1, BKT1, BLA1, BLE1, BLE2, BLH1, BLL1, BLP1, BLS1, BLV1, BMA1, BMB1, BMC1, BMH1, BNE1, BNK1, BNP1, BNT1, BPM1, BPQ1, BPT1, BPV1, BQC1, BQG1, BQH1, BQM1, BQR1, BRA1, BRD1, BSB1, BSD2, BSL2, BTH1, BTN1, BTN2, BVK1, CAE1, CCG1, CCQ1, CCV1, CDF1, CDQ1, CEQ1, CFE1, CFR1, CGL1, CGV1, CHD2, CPK1, CQD1, CQF1, CQG1, CQP1, YAB1, YBG1, YBH1, YBW1, YCH1, YCH2, YCM1 | AHP1, AIM1, AKB1, AKP1, AKV1, ALG1, ANG1, APV1, AST1, BAK1, BBI1, BEH1, BHB1, BHF1, BHM1, BHN1, BKR1, BLA1, BLE1, BLL1, BLN1, BLP1, BLV1, BMA1, BMK1, BNI1, BNK1, BPQ1, BPT1, BQG1, BQH1, BQI1, BQP1, BQQ1, BQR1, BRA1, BRB1, BRC1, BRD1, BSB1, BTH1, CAD1, CCV1, CQD1, YAR1, YBU1 |

### Slide 2
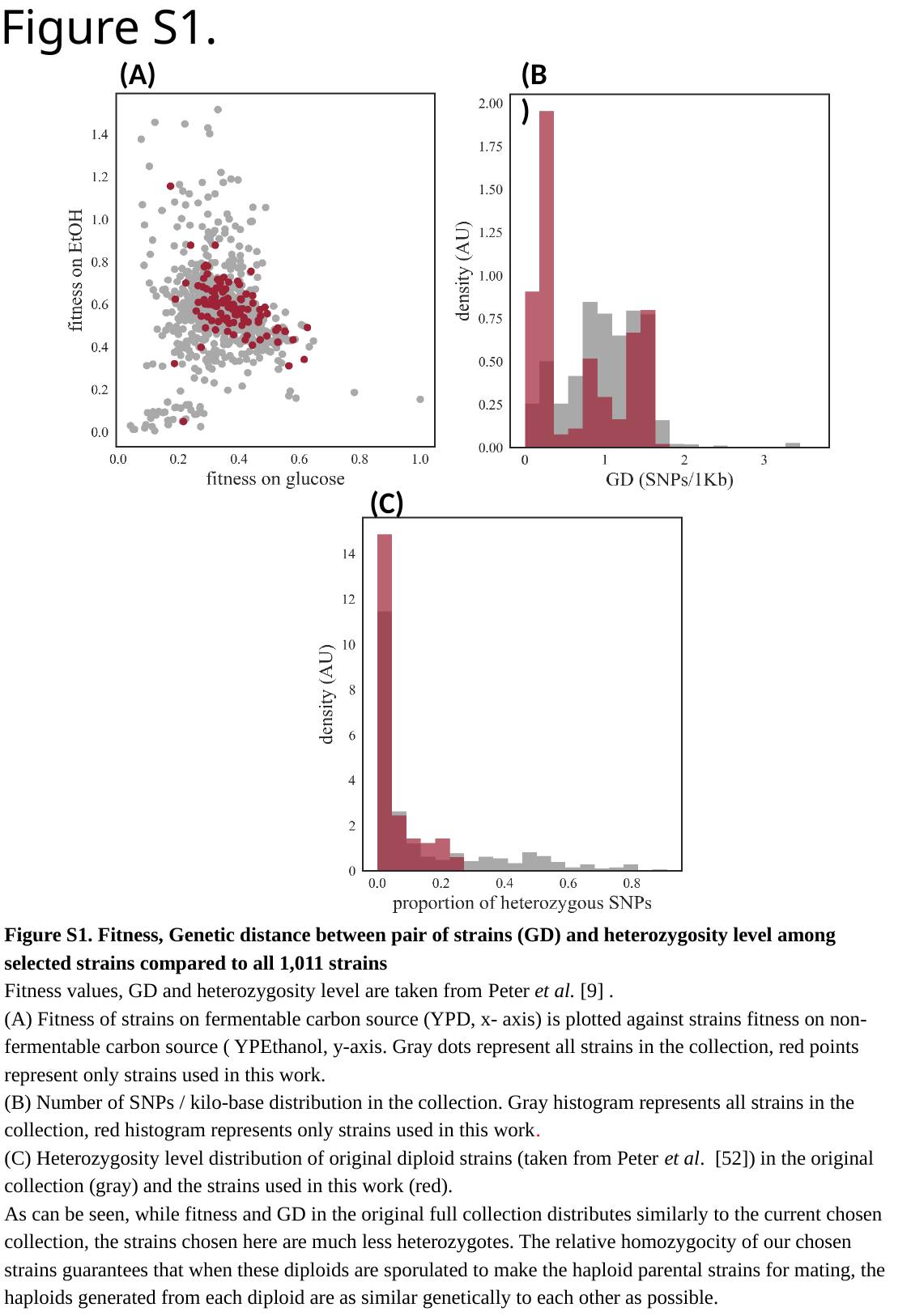

#### Figure S1.
(B)
(A)
(C)
Figure S1. Fitness, Genetic distance between pair of strains (GD) and heterozygosity level among selected strains compared to all 1,011 strains
Fitness values, GD and heterozygosity level are taken from Peter et al. [9] .
(A) Fitness of strains on fermentable carbon source (YPD, x- axis) is plotted against strains fitness on non-fermentable carbon source ( YPEthanol, y-axis. Gray dots represent all strains in the collection, red points represent only strains used in this work.
(B) Number of SNPs / kilo-base distribution in the collection. Gray histogram represents all strains in the collection, red histogram represents only strains used in this work.
(C) Heterozygosity level distribution of original diploid strains (taken from Peter et al. [52]) in the original collection (gray) and the strains used in this work (red).
As can be seen, while fitness and GD in the original full collection distributes similarly to the current chosen collection, the strains chosen here are much less heterozygotes. The relative homozygocity of our chosen strains guarantees that when these diploids are sporulated to make the haploid parental strains for mating, the haploids generated from each diploid are as similar genetically to each other as possible.

### Slide 3
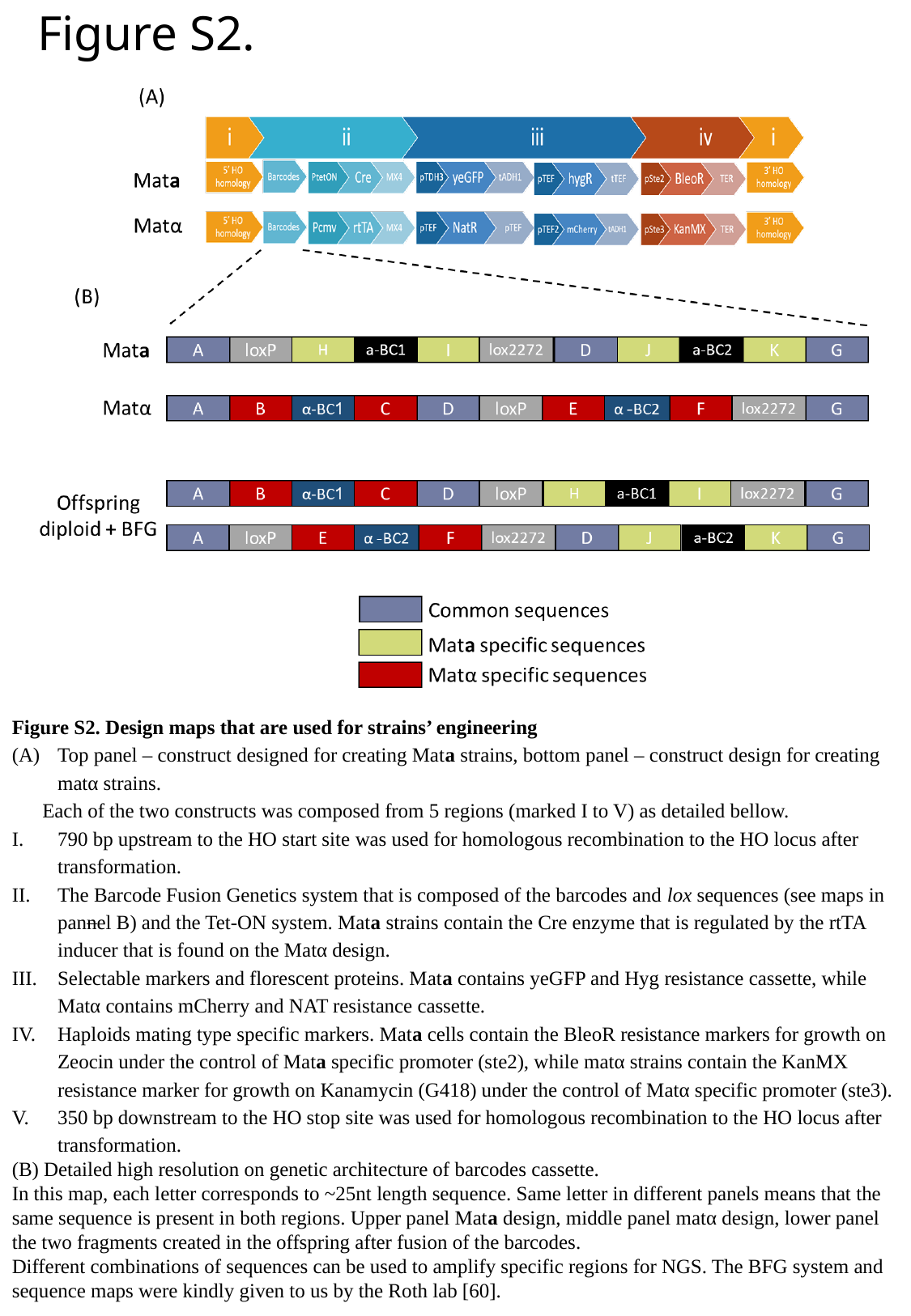

#### Figure S2.
Figure S2. Design maps that are used for strains’ engineering
Top panel – construct designed for creating Mata strains, bottom panel – construct design for creating matα strains.
Each of the two constructs was composed from 5 regions (marked I to V) as detailed bellow.
790 bp upstream to the HO start site was used for homologous recombination to the HO locus after transformation.
The Barcode Fusion Genetics system that is composed of the barcodes and lox sequences (see maps in pannel B) and the Tet-ON system. Mata strains contain the Cre enzyme that is regulated by the rtTA inducer that is found on the Matα design.
Selectable markers and florescent proteins. Mata contains yeGFP and Hyg resistance cassette, while Matα contains mCherry and NAT resistance cassette.
Haploids mating type specific markers. Mata cells contain the BleoR resistance markers for growth on Zeocin under the control of Mata specific promoter (ste2), while matα strains contain the KanMX resistance marker for growth on Kanamycin (G418) under the control of Matα specific promoter (ste3).
350 bp downstream to the HO stop site was used for homologous recombination to the HO locus after transformation.
(B) Detailed high resolution on genetic architecture of barcodes cassette.
In this map, each letter corresponds to ~25nt length sequence. Same letter in different panels means that the same sequence is present in both regions. Upper panel Mata design, middle panel matα design, lower panel the two fragments created in the offspring after fusion of the barcodes.
Different combinations of sequences can be used to amplify specific regions for NGS. The BFG system and sequence maps were kindly given to us by the Roth lab [60].

### Slide 4
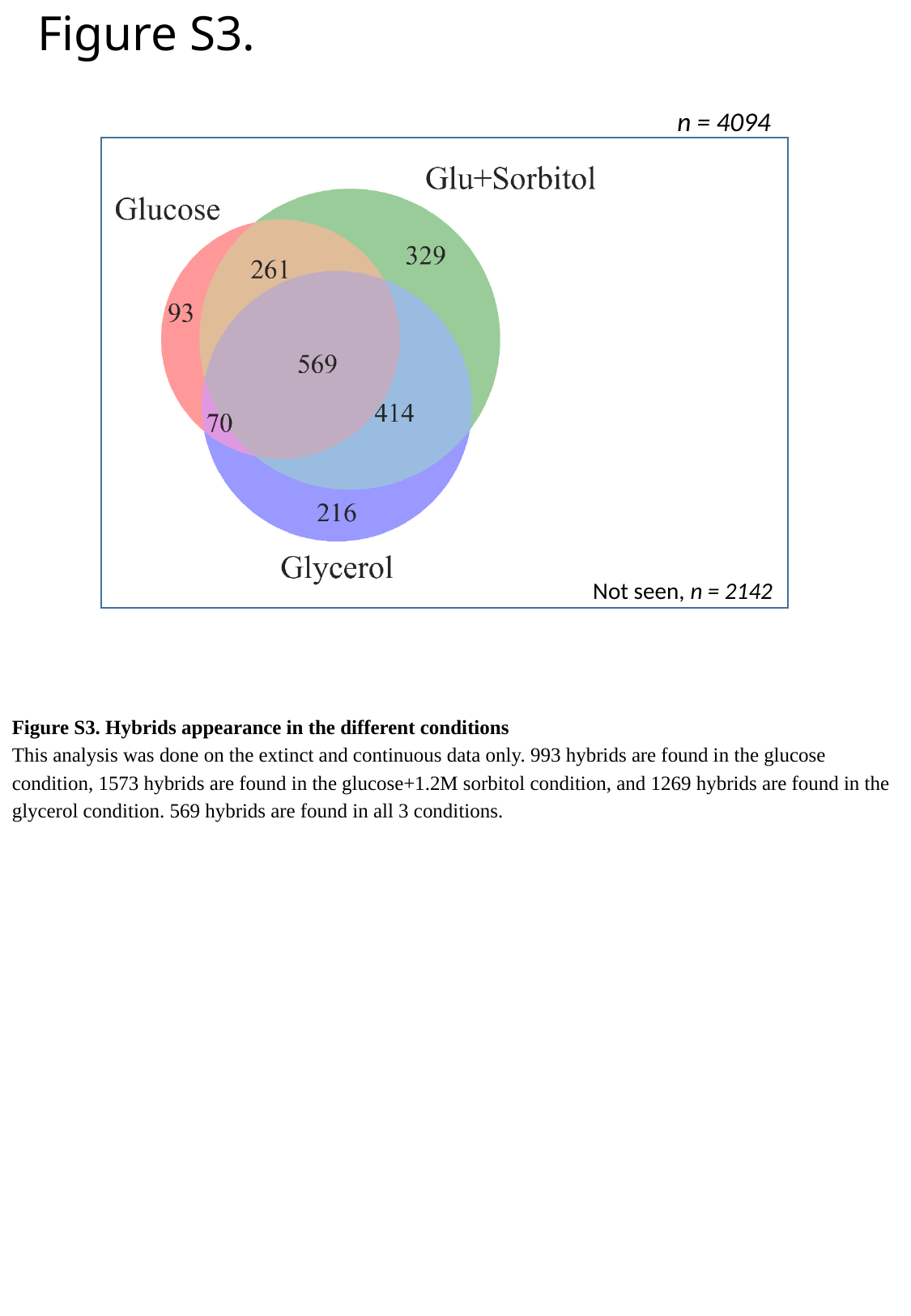

Figure S3.
n = 4094
Not seen, n = 2142
Figure S3. Hybrids appearance in the different conditions
This analysis was done on the extinct and continuous data only. 993 hybrids are found in the glucose condition, 1573 hybrids are found in the glucose+1.2M sorbitol condition, and 1269 hybrids are found in the glycerol condition. 569 hybrids are found in all 3 conditions.

### Slide 5
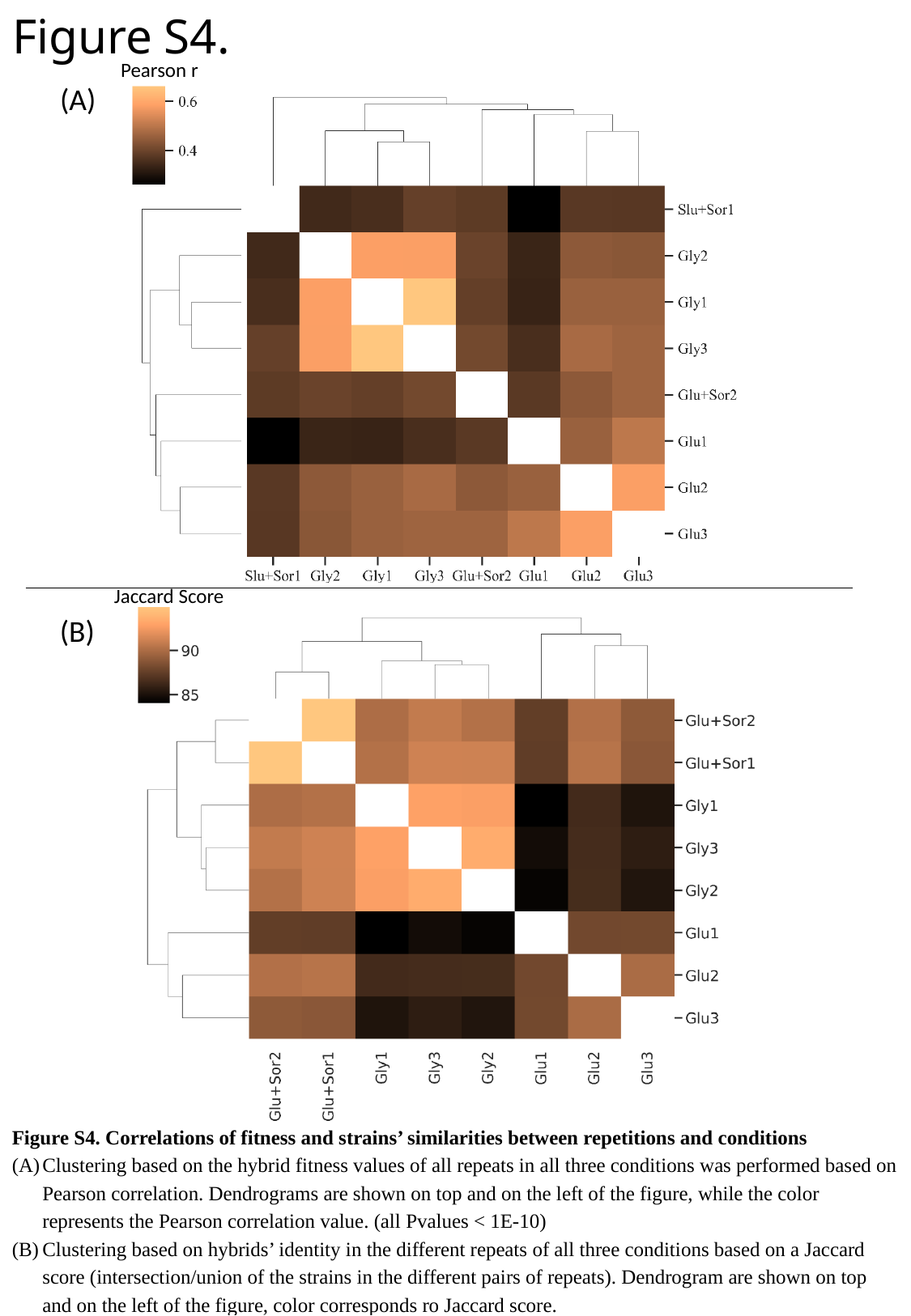

Figure S4.
Pearson r
(A)
Jaccard Score
(B)
Figure S4. Correlations of fitness and strains’ similarities between repetitions and conditions
Clustering based on the hybrid fitness values of all repeats in all three conditions was performed based on Pearson correlation. Dendrograms are shown on top and on the left of the figure, while the color represents the Pearson correlation value. (all Pvalues < 1E-10)
Clustering based on hybrids’ identity in the different repeats of all three conditions based on a Jaccard score (intersection/union of the strains in the different pairs of repeats). Dendrogram are shown on top and on the left of the figure, color corresponds ro Jaccard score.

### Slide 6
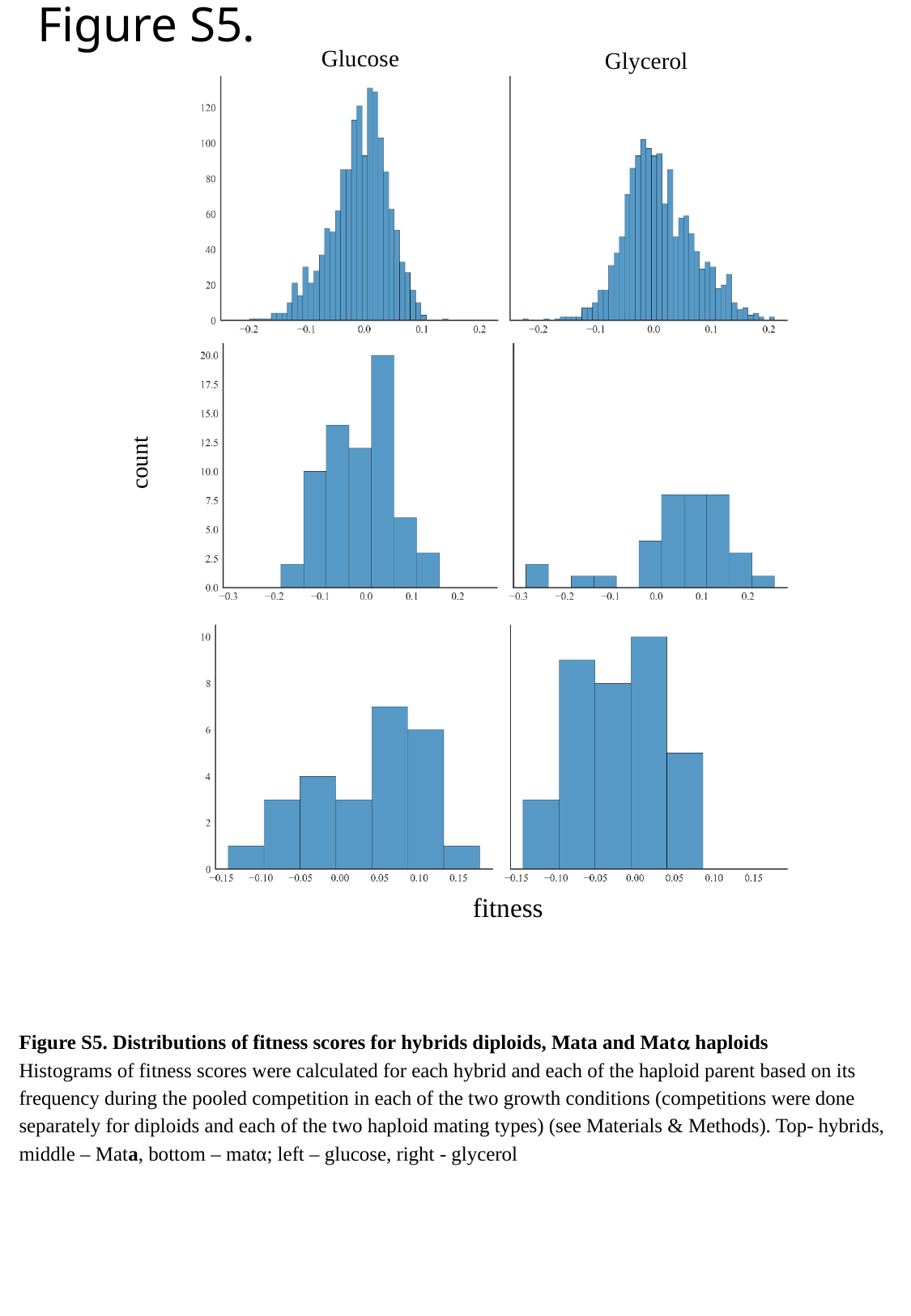

Figure S5.
Glucose
Glycerol
count
fitness
Figure S5. Distributions of fitness scores for hybrids diploids, Mata and Matα haploids
Histograms of fitness scores were calculated for each hybrid and each of the haploid parent based on its frequency during the pooled competition in each of the two growth conditions (competitions were done separately for diploids and each of the two haploid mating types) (see Materials & Methods). Top- hybrids, middle – Mata, bottom – matα; left – glucose, right - glycerol

### Slide 7
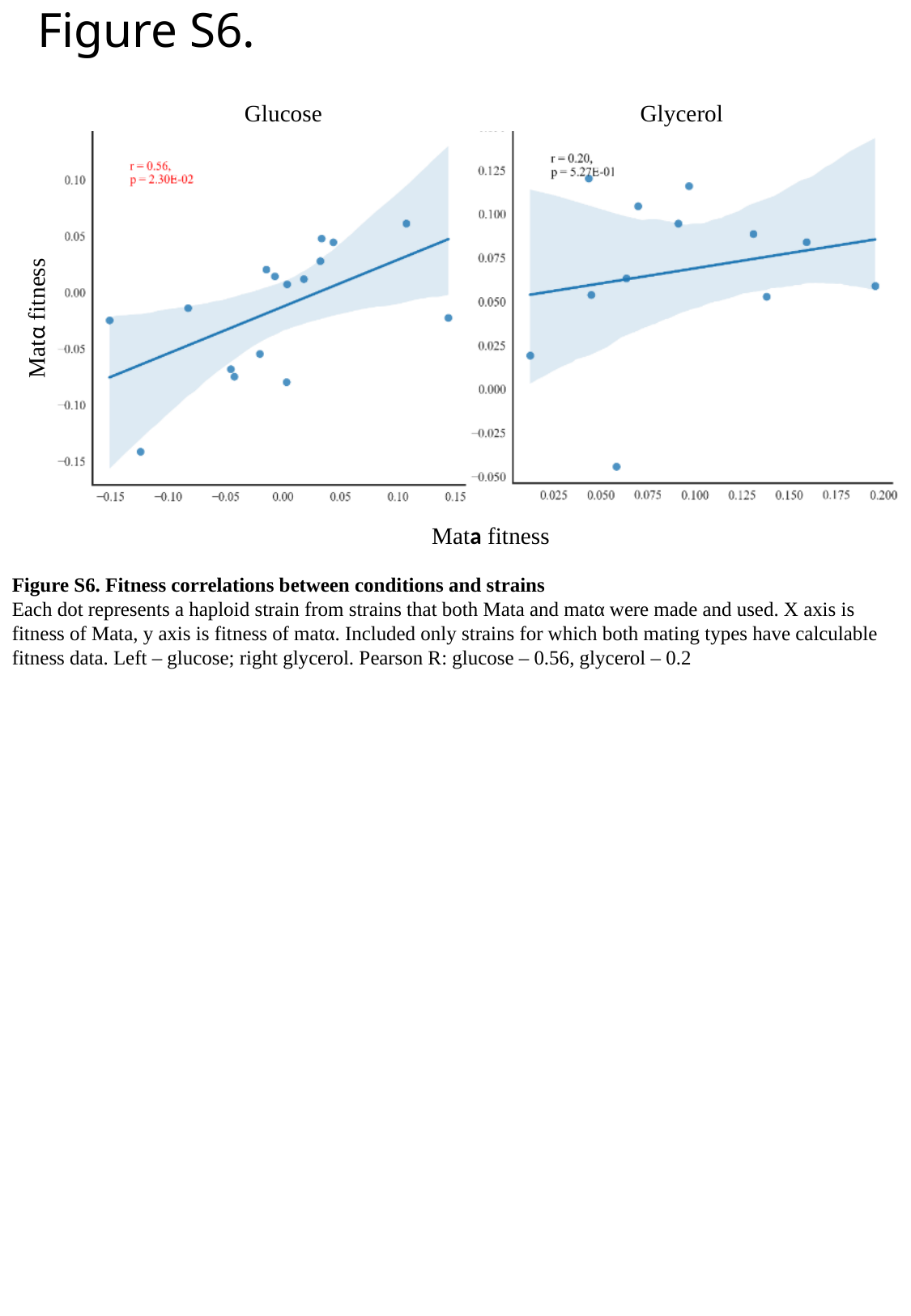

Figure S6.
Glycerol
Glucose
Matα fitness
Mata fitness
Figure S6. Fitness correlations between conditions and strains
Each dot represents a haploid strain from strains that both Mata and matα were made and used. X axis is fitness of Mata, y axis is fitness of matα. Included only strains for which both mating types have calculable fitness data. Left – glucose; right glycerol. Pearson R: glucose – 0.56, glycerol – 0.2

### Slide 8
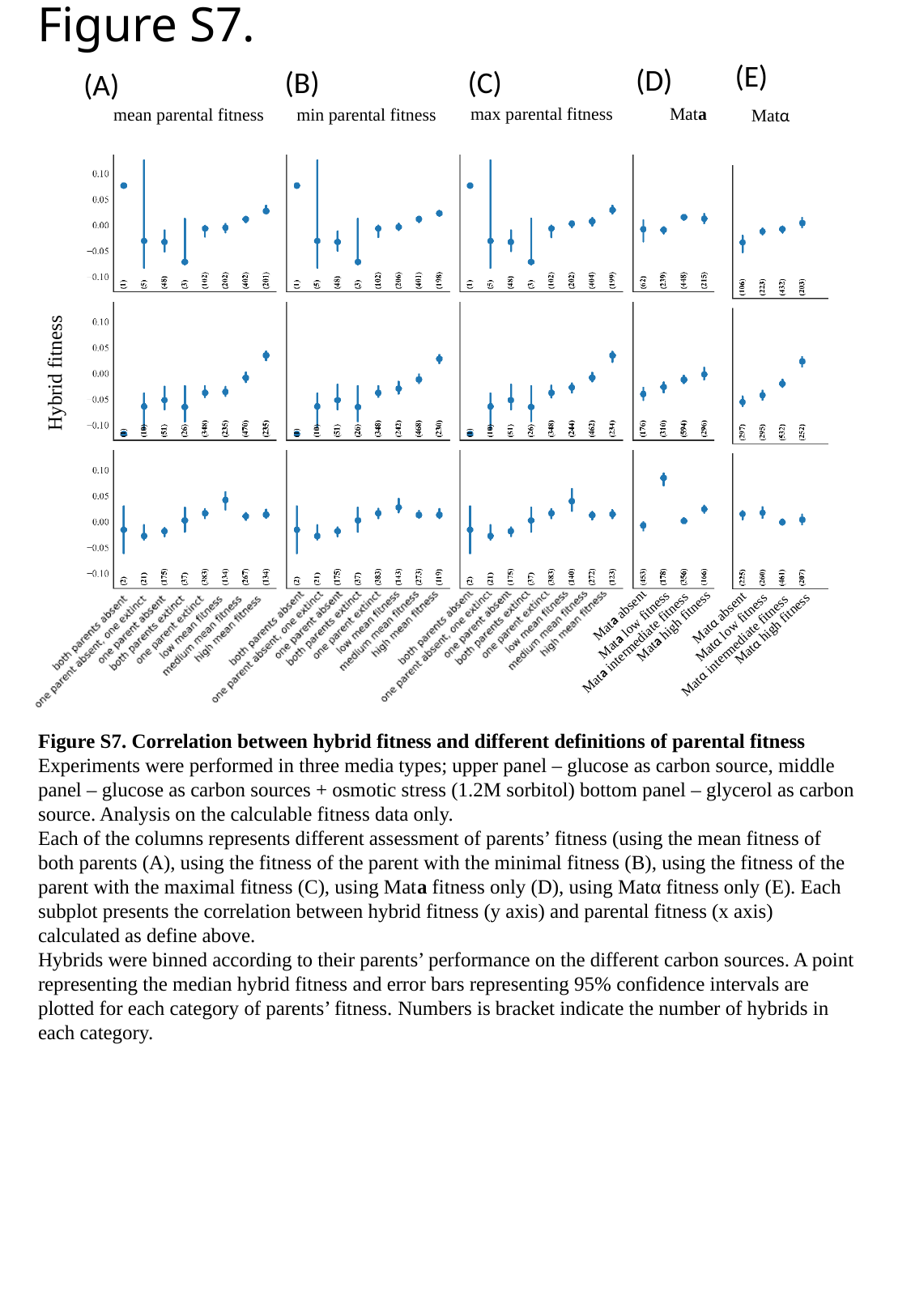

Figure S7.
(E)
(D)
(B)
(C)
(A)
Mata
max parental fitness
mean parental fitness
min parental fitness
Matα
Hybrid fitness
Mata absent
Matα absent
Mata low fitness
Mata high fitness
Matα low fitness
Matα high fitness
Mata intermediate fitness
Matα intermediate fitness
Figure S7. Correlation between hybrid fitness and different definitions of parental fitness
Experiments were performed in three media types; upper panel – glucose as carbon source, middle panel – glucose as carbon sources + osmotic stress (1.2M sorbitol) bottom panel – glycerol as carbon source. Analysis on the calculable fitness data only.
Each of the columns represents different assessment of parents’ fitness (using the mean fitness of both parents (A), using the fitness of the parent with the minimal fitness (B), using the fitness of the parent with the maximal fitness (C), using Mata fitness only (D), using Matα fitness only (E). Each subplot presents the correlation between hybrid fitness (y axis) and parental fitness (x axis) calculated as define above.
Hybrids were binned according to their parents’ performance on the different carbon sources. A point representing the median hybrid fitness and error bars representing 95% confidence intervals are plotted for each category of parents’ fitness. Numbers is bracket indicate the number of hybrids in each category.

### Slide 9
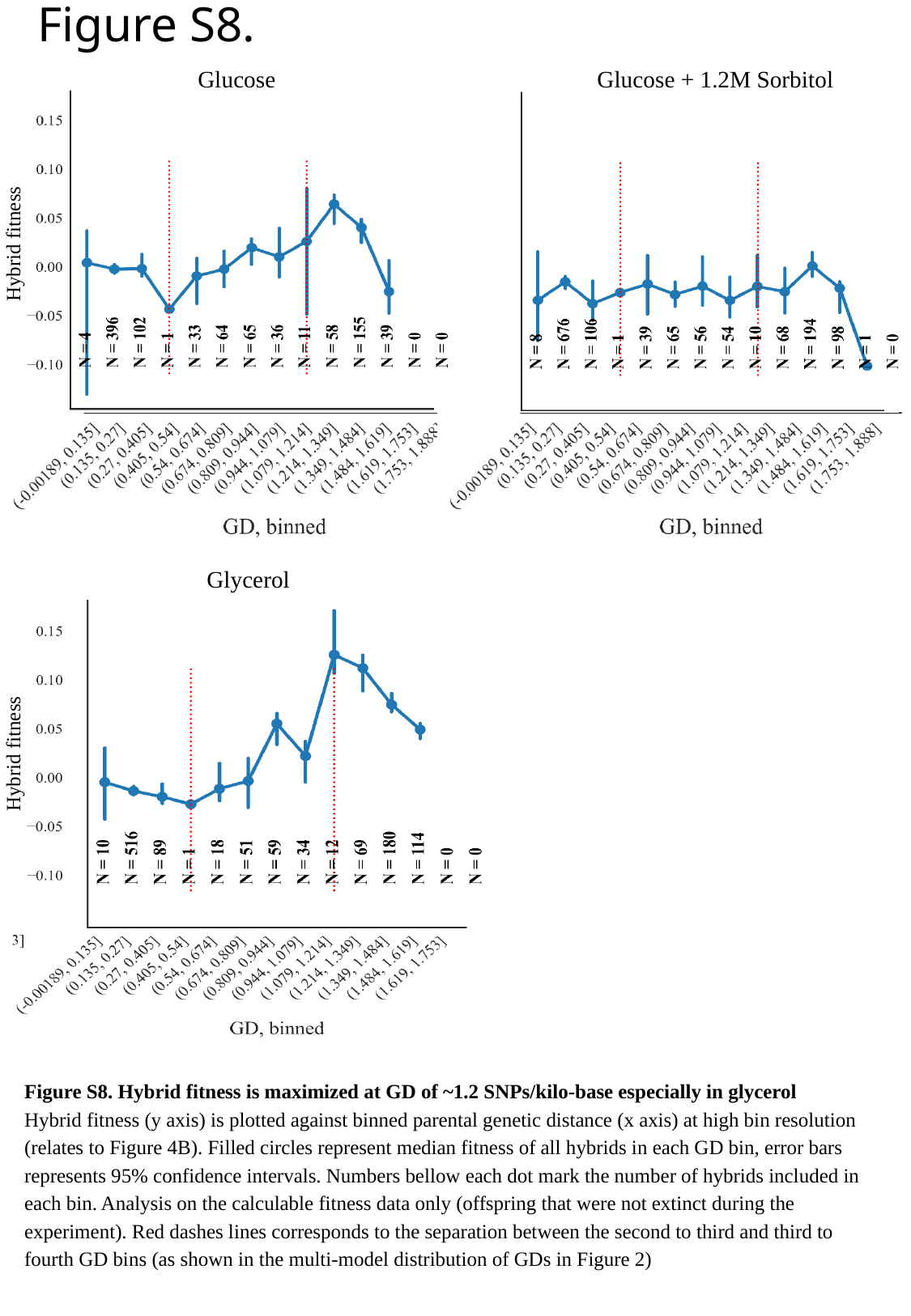

Figure S8.
Glucose
Glucose + 1.2M Sorbitol
Hybrid fitness
Glycerol
Hybrid fitness
Figure S8. Hybrid fitness is maximized at GD of ~1.2 SNPs/kilo-base especially in glycerol
Hybrid fitness (y axis) is plotted against binned parental genetic distance (x axis) at high bin resolution (relates to Figure 4B). Filled circles represent median fitness of all hybrids in each GD bin, error bars represents 95% confidence intervals. Numbers bellow each dot mark the number of hybrids included in each bin. Analysis on the calculable fitness data only (offspring that were not extinct during the experiment). Red dashes lines corresponds to the separation between the second to third and third to fourth GD bins (as shown in the multi-model distribution of GDs in Figure 2)

### Slide 10
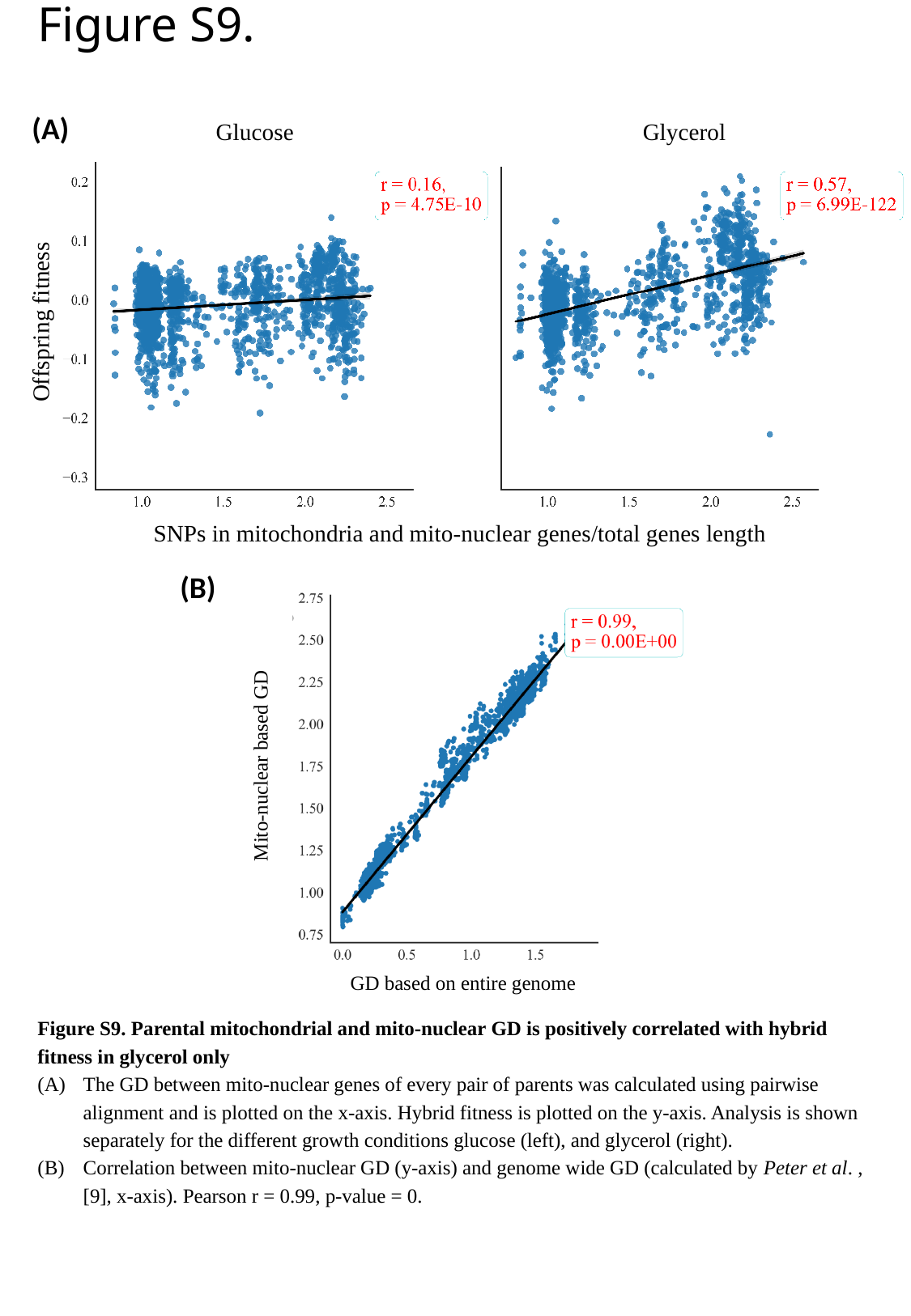

Figure S9.
(A)
Glucose
Glycerol
Offspring fitness
SNPs in mitochondria and mito-nuclear genes/total genes length
(B)
Mito-nuclear based GD
GD based on entire genome
Figure S9. Parental mitochondrial and mito-nuclear GD is positively correlated with hybrid fitness in glycerol only
The GD between mito-nuclear genes of every pair of parents was calculated using pairwise alignment and is plotted on the x-axis. Hybrid fitness is plotted on the y-axis. Analysis is shown separately for the different growth conditions glucose (left), and glycerol (right).
Correlation between mito-nuclear GD (y-axis) and genome wide GD (calculated by Peter et al. , [9], x-axis). Pearson r = 0.99, p-value = 0.

### Slide 11
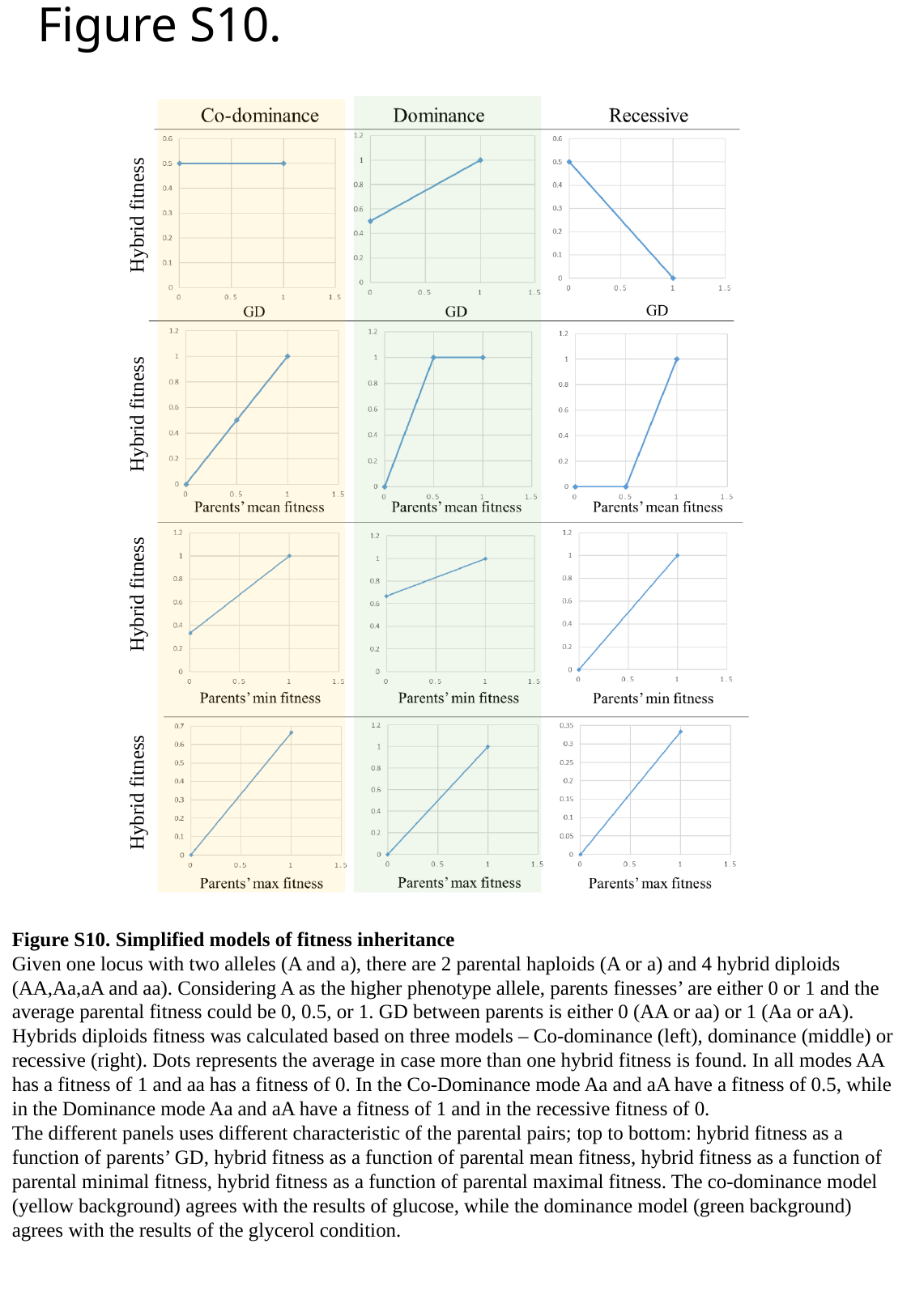

Figure S10.
Hybrid fitness
Hybrid fitness
Hybrid fitness
Hybrid fitness
Figure S10. Simplified models of fitness inheritance
Given one locus with two alleles (A and a), there are 2 parental haploids (A or a) and 4 hybrid diploids (AA,Aa,aA and aa). Considering A as the higher phenotype allele, parents finesses’ are either 0 or 1 and the average parental fitness could be 0, 0.5, or 1. GD between parents is either 0 (AA or aa) or 1 (Aa or aA). Hybrids diploids fitness was calculated based on three models – Co-dominance (left), dominance (middle) or recessive (right). Dots represents the average in case more than one hybrid fitness is found. In all modes AA has a fitness of 1 and aa has a fitness of 0. In the Co-Dominance mode Aa and aA have a fitness of 0.5, while in the Dominance mode Aa and aA have a fitness of 1 and in the recessive fitness of 0.
The different panels uses different characteristic of the parental pairs; top to bottom: hybrid fitness as a function of parents’ GD, hybrid fitness as a function of parental mean fitness, hybrid fitness as a function of parental minimal fitness, hybrid fitness as a function of parental maximal fitness. The co-dominance model (yellow background) agrees with the results of glucose, while the dominance model (green background) agrees with the results of the glycerol condition.

### Slide 12
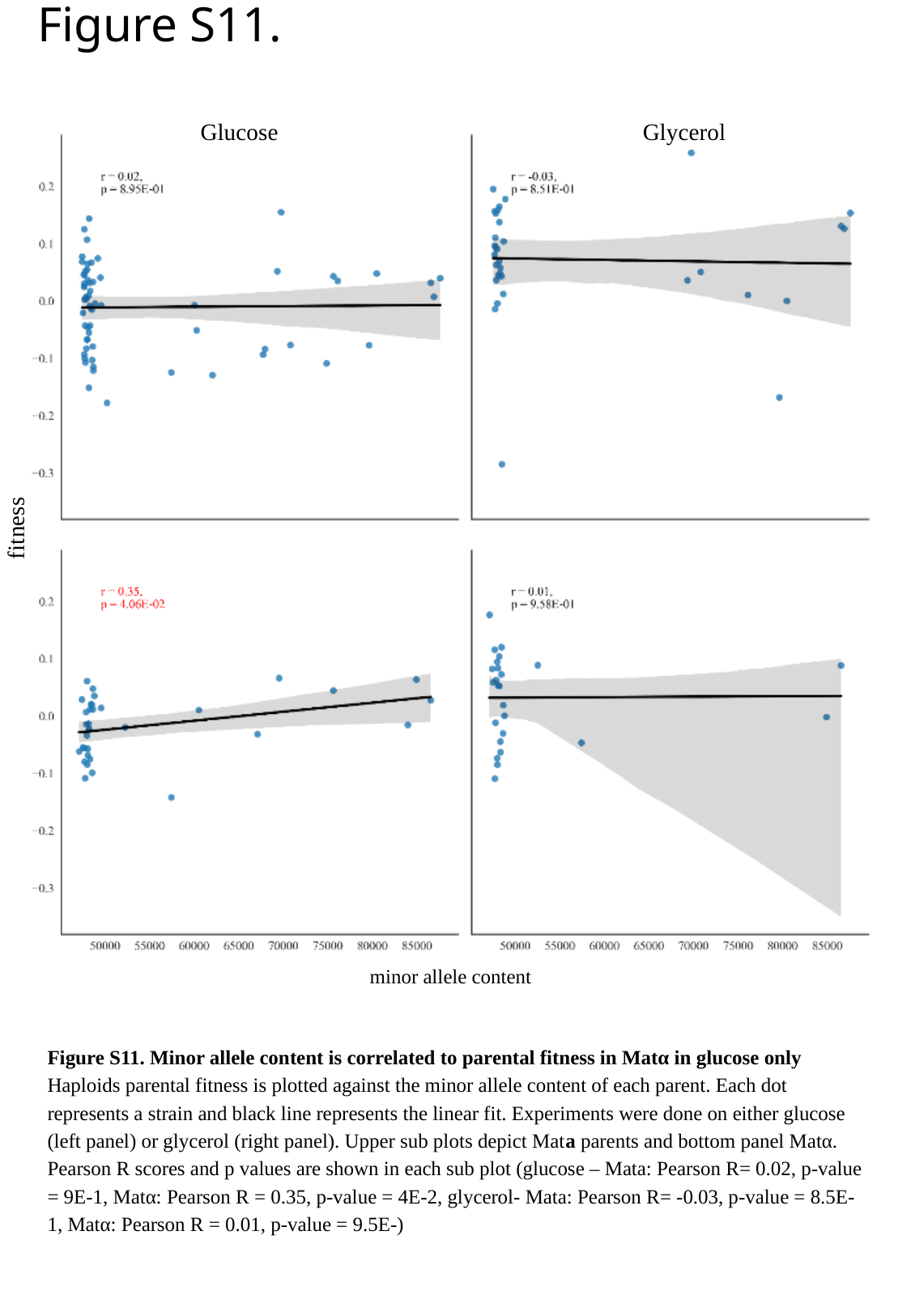

Figure S11.
Glucose
Glycerol
fitness
minor allele content
Figure S11. Minor allele content is correlated to parental fitness in Matα in glucose only
Haploids parental fitness is plotted against the minor allele content of each parent. Each dot represents a strain and black line represents the linear fit. Experiments were done on either glucose (left panel) or glycerol (right panel). Upper sub plots depict Mata parents and bottom panel Matα. Pearson R scores and p values are shown in each sub plot (glucose – Mata: Pearson R= 0.02, p-value = 9E-1, Matα: Pearson R = 0.35, p-value = 4E-2, glycerol- Mata: Pearson R= -0.03, p-value = 8.5E-1, Matα: Pearson R = 0.01, p-value = 9.5E-)
