## Supplementary Note 1 for "Quantitative genetics of natural *S. cerevisiae* strains upon sexual mating reveals heritable determinants of cellular fitness"

### Supplementary note 1**: *An experimental platform to study fitness inheritance in yeast on a massive scale***

###### Strains choice

Strains were chosen from a collection of ~1000 *S. cerevisiae* strains isolated from diverse ecological niches all over the world (described in Peter *et al*. (Peter *et al.*, 2018)). From those, 200 strains were chosen for genetic engineering (**Figure 1A** and **Figure S1**). Strains were chosen based on the following parameters; Euploid diploids, intact HO and sporulation rate were all chosen to allow easier manipulation of strains during the genetic engineering steps. Euploid diploids were chosen to reduce difficulties of mating efficiency; an intact HO was chosen to allow transformation into the HO in the genetic engineering part. Efficient sporulation is needed to allow sporulation after transformation. In addition to the technical criteria above, three parameters were optimized to allow enough statistical power to address the roles of GD and parents’ fitness on hybrids. Yeast usually have two opposing metabolic preferences for energy extraction, fermentation or respiration. While yeast that live on glucose ferments, different carbon sources such as glycerol or ethanol cause yeast to respirate aerobically. Using Peter *et al.* data (Peter *et al*., 2018), we observed a negative correlation between strains fitness in glucose (YPD) and ethanol (YPEthanol) (shown in **Figure S1A**). Strains were chosen to represent the following groups, either high fitness on glucose and low on ethanol, high fitness on ethanol and low on glucose or intermediate fitness on both. Few strains with low fitness on both media were also added. A second criterion for strain choice was low heterozygosity of the strains. The genetic distance between strains was determined by the data of Peter *et al.* and as strains had to be sporulated in the construction process, high heterozygosity strains produce haploid cells that differ from the original strains and thus deflect the genetic distance matrix (**Figure S1C**). In addition, strains were chosen to represent the original distribution of GD as in the entire collection (**Figure S1B)**. As explained in the introduction, organisms GD may influence hybrid fitness (such in the case of hybrid vigor), thus, we wanted to explore the connection between GD and fitness in different carbon sources. To that end, it was important to have as many genetic distances as possible between pairs of strains. Furthermore, strains were chosen such that they can be binned in four bins: very low (“selfing”), low, intermediate and high GD to allow different statistical analysis methods (**Figure 2B**).

###### Strains’ construction

Massive genetic engineering was done to be able to work with natural isolates in this complex and high throughput experiments. As original strains were natural isolates that can typically switch in the haploid form between mating types, knock out of the HO locus was done to disable mating type switching (Haber, 2012). As the experiments involved selection of either haploids or diploids cells, as well as easily detecting haploids and diploids in FACS analyzer, strains were transformed with constitutive antibiotics cassettes and fluorescence markers (Mat**a**: HygR-GFP, Matα: NatR-mCherry). “Magic Markers” (adapted from (Tong & Boone, 2007)) were also added, enabling the selection of a specific mating type after sporulation. Lastly, a sophisticated system enabling fusion of barcodes originating from different chromosomes was added as well to allow identification of hybrids after *en masse* mating. Below is an explanation of each of the above requirements. The full construct is shown in **Figure S2** and it includes the following regions: (i) genome homology region: 790bp and 350bp homology (5’ and 3’ respectively) to the HO locus on both ends of the construct (**Figure S2A**). The specific sequences of homology were chosen to have the least amount of SNPs between most strains, thus enabling favorable conditions for genome integration. Integration to the HO and by that knocking it out is essential to achieve stable haploids, as needed for further steps in the project (ii) Barcode Fusion Genetic (BFG): This region was kindly given to us by the lab of Fredrick Roth (Yachie *et al*., 2016). This region enables the identification of the two parents of the hybrids using Next Generation Sequencing (NGS). Each parental strain is labeled with two barcodes, flanked by a loxP/lox2272 sites. Activating Cre recombinase in the hybrids diploid (with tetracycline antibiotic), fuses the parental barcodes, resulting in one linear fragments containing one barcode originating from parent Mat**a** and one barcode of parent Matα that could be identified using NGS (Also see **Figure S2B**) (iii) Constitutive markers and fluorescence proteins. Constitutive markers enable the selection of diploids after transformation. Diploids that received the Mat**a** design are selected using Hygromycin (Hyg) and further validated by FACS for the presence of GFP. Diploids that received the Matα design are selected using Nourseothricin (Nat) and further validated by FACS for the presence of mCherry. (iv) Haploid selecting markers. After transformation, the strains are sporulated, and haploids that contain either Mat**a** or Matα designs are selected by Zeocin or Geneticin (G418), respectively. Since resistance cassettes are under the control of mating type specific promoters; Ste2 promoter is active in Mat**a** cells only, while Ste3 promoter is active in Matα cells only, this part of the construct allows the selection of the desired haploids.

To achieve final engineered strains, strains were transformed, selected on antibiotic and verified by FACS, and then sporulated. After sporulation, haploid strains were selected and eventually verified by Sanger sequencing. At the end of this process from 200 initial diploid strains chosen as described in the section “strain choice” 89 Mat**a** engineered strains and 46 Matα engineered strains were obtained. The verified final strain set has similar properties to the chosen strain set, without any dramatic effect on the collection fitness, GD or heterozygosity

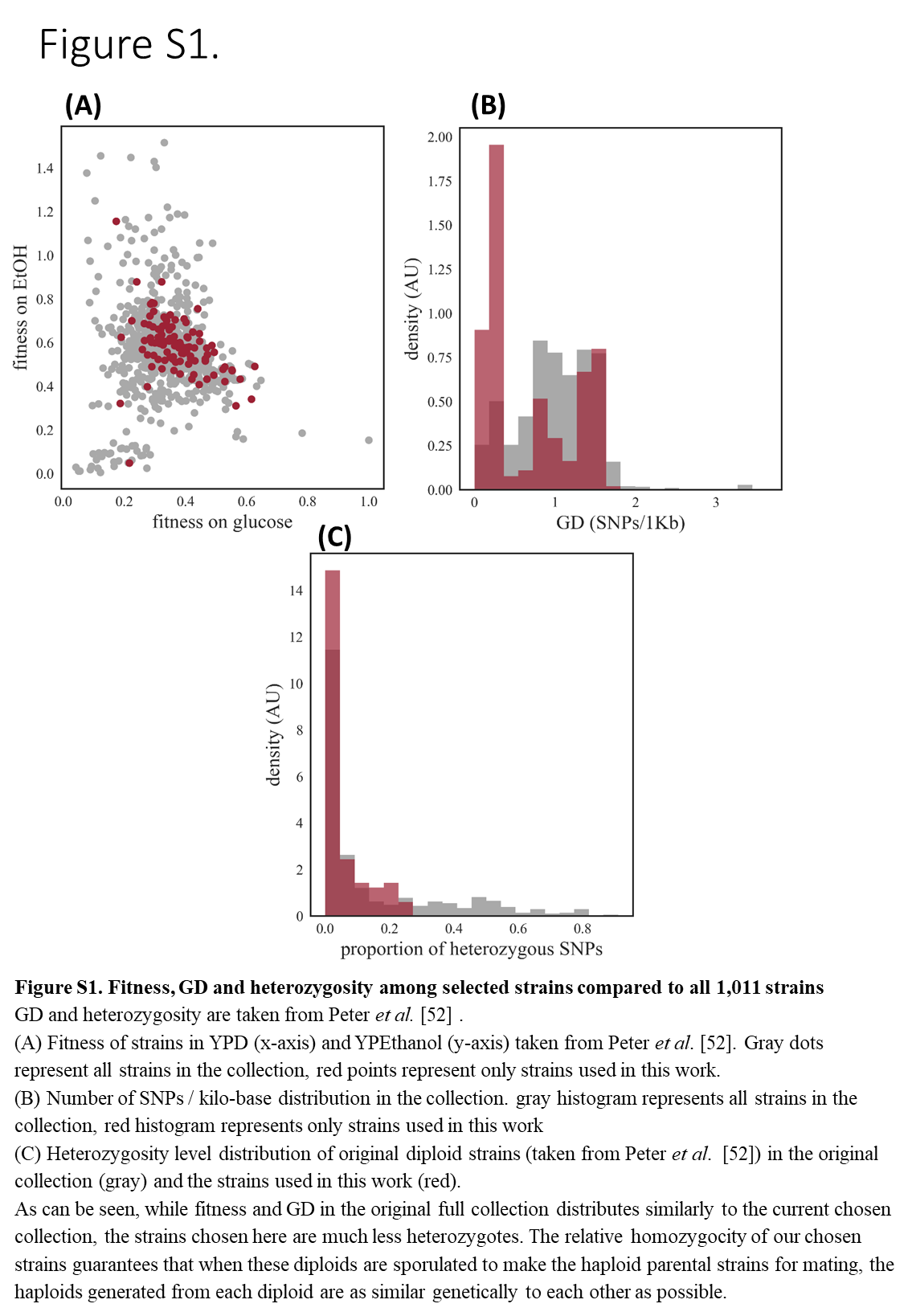

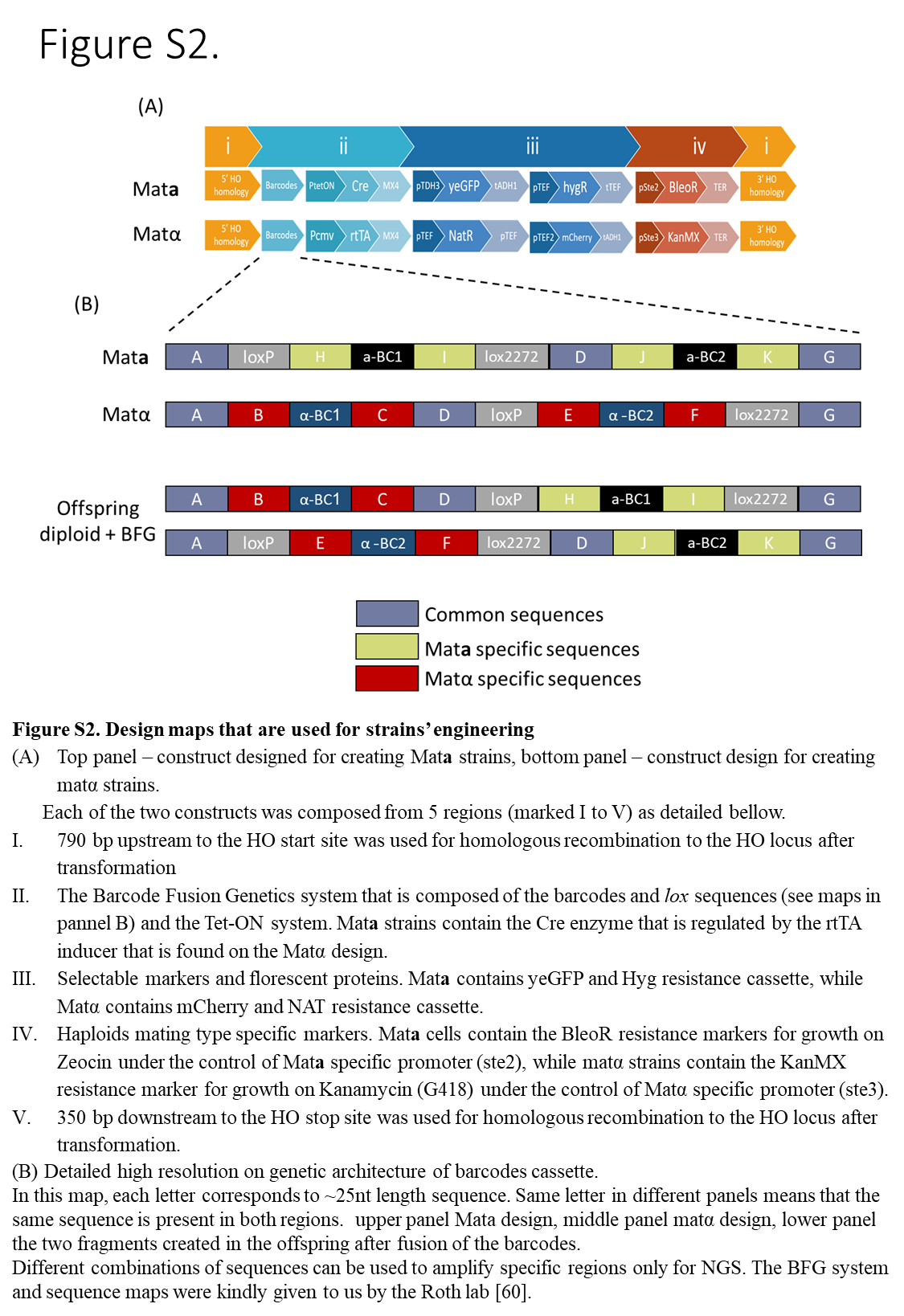
