## Supplementary Note 2 for "Quantitative genetics of natural *S. cerevisiae* strains upon sexual mating reveals heritable determinants of cellular fitness"

*Dominance and Co-dominance modes of inheritance explain the inheritance of fitness in glycerol and glucose, respectively*

We hypothesized that an optional explanation for the differences in fitness inheritance in the two carbon sources may be due to different genes with different modes of inheritance that are involved in affecting fitness in these conditions. To shed more light on this hypothesis, we constructed a simple theoretical model of fitness inheritance that implements different inheritance modes, in similarity to [41]. In particular, we examined cases of dominance (i.e., when the heterozygote is as fit as the homozygote of the better allele), recessive (i.e., when the heterozygote is fit as the homozygote of the worse allele, “aa”), and co-dominance (i.e., the heterozygote has a fitness equal to the mean fitness of the two homozygote types). Under each of these inheritance models we examined how fitness of hybrids depends on parental fitness and the GD between them. Of all these models, the dominance model generates behavior similar to the results observed in glycerol, while the co-dominance model fits better the observations in glucose (**Figure S10**). In particular, the co-dominance (yellow background column in **Figure S10**) model shows a positive correlation between hybrid fitness and parents’ fitness (measured either as mean, min, or max parental fitness), and also no correlation between hybrid fitness and parents’ GD, as observed in glucose. On the other hand, the dominance model (green background column in **Figure S10**) shows an increase in hybrid fitness with parents’ mean fitness at the low parental fitness range, and then a plateau of no correlation; but a positive correlation between hybrid fitness and parents’ GD, as observed in glycerol. The fit between the model and the experimental results suggests a distinct mode of inheritance in each of the conditions, and in particular an abundance of dominant inheritance loci in glycerol and co-dominance in glucose.

*A toy model fitness inheritance under different inheritance modes*

We consider a hypothetical single SNP position with two possible alleles, marked as “A” or “a”. Starting from parental haploids, of the two mating types Mata or Matα, the fitness at this SNP locus was assigned to be 1 in case of the “A” allele and 0 in the case of the “a” allele. The four possible hybrids diploids combinations are “AA”, “Aa”, “aA” and “aa”. Genetic distance between parental alleles, GD, is defined to be 0 when both parents have the same allele and 1 otherwise. Parental average fitness was calculated as the average of parents’ fitness. Likewise, parental min and max fitness are defined respectively as the fitness of the parent with the minimum or maximum fitness value. Fitness of the hybrids was calculated based on several alternative models: the dominance (heterozygous is as fit as the homozygous “A” variant) or the co-dominance (heterozygous is half as fit as the homozygous “A” variant) or the recessive (heterozygous is as fit as the homozygous “a” variant) (**Table S2**). With this model, we could plot in Figure 5 the expected dependence of hybrid fitness on parental GD and on parental fitness under each model.

Table S2. Fitness and GD values of parents and hybrids based on different inheritance models

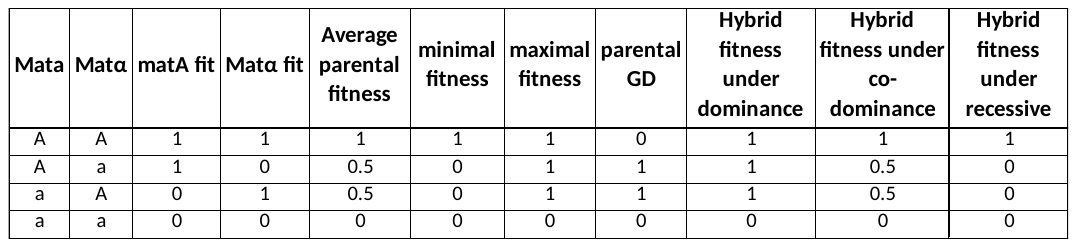

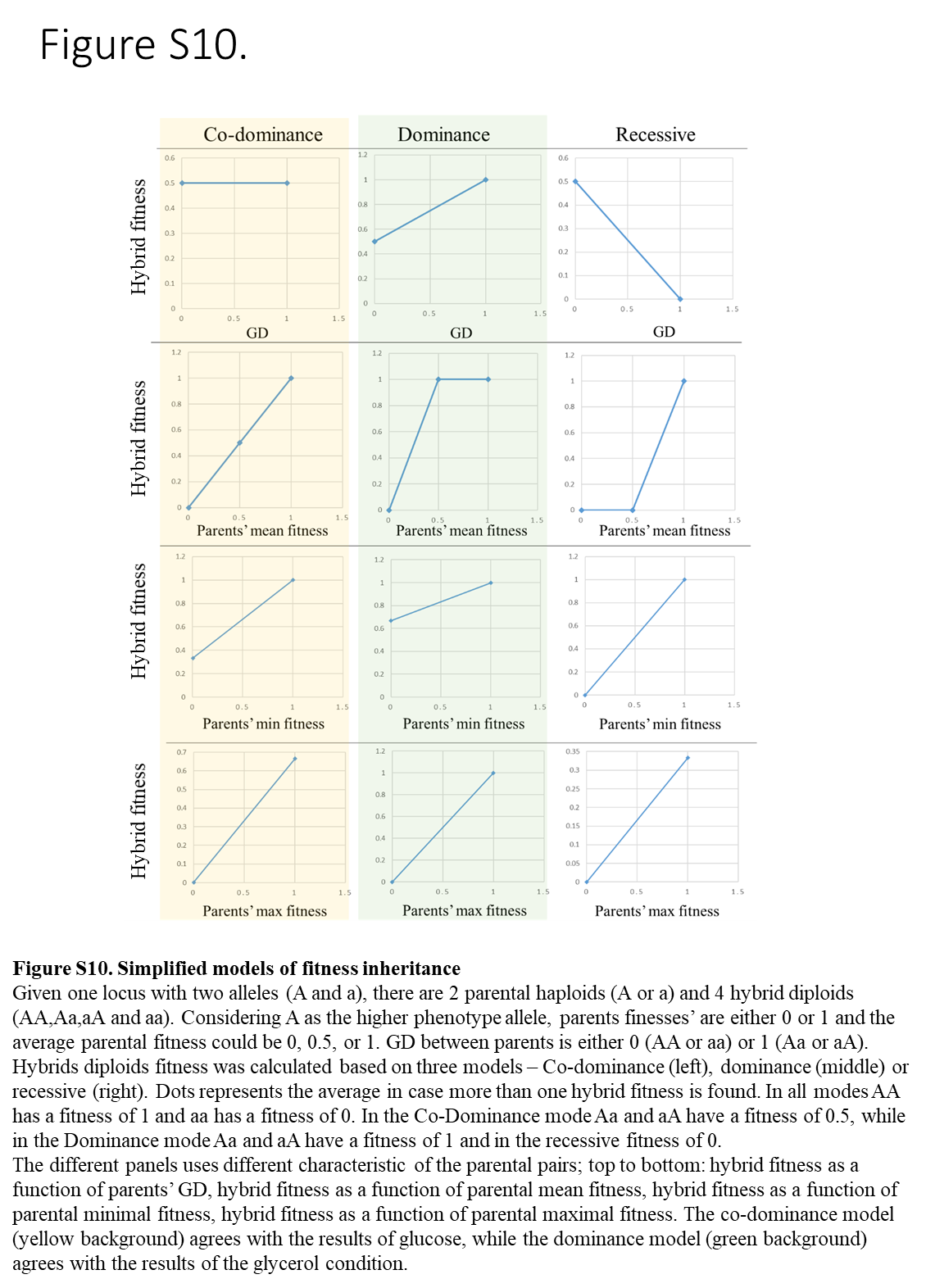
